# Improving All-Atom Molecular Dynamics Models for Quantitative Prediction of Nanopore Blockade Current

**DOI:** 10.64898/2026.06.12.731905

**Authors:** Jingqian Liu, Crystal Rodriguez, Min Chen, Aleksei Aksimentiev

## Abstract

All-atom molecular dynamics has become an indispensable tool in development of nanopore sensors of biological information. In a typical nanopore experiment, measurements of ionic current flowing through a nanopore report on the chemical structure of biomolecules that pass through the nanopore. Such experiments alone are often insufficient to relate the structure of the biomolecules to the ionic current modulations. The molecular dynamics method can establish such a relationship directly through a brute force simulation under applied electric field. Here, we examine the ability of molecular dynamics force fields to reproduce experimentally measured nanopore blockade currents produced by single-stranded DNA. Our simulations show that none of the “off the shelf” force fields (CHARMM36, AMBER Parmbsc1 and DES-AMBER) is capable of reproducing experimental data with the desired level of accuracy. To improve the accuracy, we examined and refined interactions between ions, protein nanopores and DNA, guided by experiments designed specifically for this purpose. Ultimately, the introduction of surgical corrections to non-bonded interactions within the CHARMM36 force field produced a favorable agreement between simulation and experiment. This refined parameterization, initially developed for nanopore sensing simulations, may have broader applications in computational studies of DNA–protein systems.

## INTRODUCTION

Nanopore sequencing has revolutionized nucleic acid characterization, enabling real-time, label-free analysis of long DNA or RNA molecules. ^1–3^ In a typical measurement,^4,5^ a biological nanopore is embedded in a lipid bilayer, and a nucleic acid strand is electrophoretically threaded through it. A DNA motor enzyme advances the strand one nucleotide at a time, while an applied transmembrane voltage drives ionic current through the pore. As each nucleotide passes through the sensing region of the nanopore, it modulates the ionic current in a nucleotide type-specific manner. This stepwise current blockade is then transformed into the nucleotide sequence of the passing nucleic acid strand using a deconvolution algorithm ^6^ trained on a sequence–current lookup table.^7^

Although the first nanopore experiments were done using a biological nanopore alpha-hemolysin,^8–10^ the vast majority of DNA or RNA sequencing experiments are currently done using either MspA^11^ or CsgG^12^ nanopores, heavily modified from their naturally occurring forms to increase their stability and nucleotide recognition. Nanopore sequencing using CsgG was successfully commercialized by the Oxford Nanopore Technologies, Inc. to provide portable and high-throughput sequencing platforms.^13^ CsgG is prized for its robust performance in single-molecule sequencing, compatibility with motor enzymes, and ability to accommodate long nucleic acid strands, making nanopore sequencing suitable for real-world applications.^14,15^ In contrast, the MspA nanopore is more frequently used in academic research laboratories.^16–20^ Most notably, it was the first nanopore to decode the nucleotide sequence of a natural DNA polymer,^7^ paving the way for commercial nanopore sequencing. MspA has a narrow constriction^11^ that is particularly well-matched to the diameter of single-stranded DNA (ssDNA), which provides high contrast for nucleotide identification.^21^ In addition, MspA is structurally stable, genetically tractable, and readily available, making it a popular platform for probing the physical principles of nanopore sensing^22–24^ and testing new analytical methods.^25–27^

Building on protocols developed for MD simulations of membrane channels, ^28^ MD simulations of nanopore transport began from determining, *in silico*, the current–voltage dependence of alpha-hemolysin.^29^ The simulations revealed strikingly good agreement between experiment and simulations, which not only reproduced the magnitude and direction of current rectification but also the absolute value of the current at experimental voltages. The first simulations of ssDNA transport,^30^ however, resolved to quantitative analysis of ssDNA conformations under nanopore confinement, which nevertheless was sufficient to explain experimentally observed directional asymmetry of the transport. The principle difficulty precluding direct comparison of simulation and experiment was the orders-of-magnitude mismatch in the time scale of the MD method (*< µ*s) and the time scale of ssDNA translocation (tens of milliseconds).

Several methods have been developed to overcome the time scale problem, including grid-steered MD,^31^ multicopy simulations of minimal nanopore models^32^ and the use of a special purpose supercomputer, Anton.^33^ Using the first generation of Anton machines (Anton 1), long (tens of *µ*s) continuous trajectories of ssDNA behavior in MspA were obtained. ^34^ The simulations revealed major fluctuations of ionic current at the microsecond time scale caused by the stochastic motion of ssDNA strands within the nanopore. Despite the fluctuations, the simulations identified the number of bulk-like water molecules in the nanopore constriction, *i.e.,* its conductive volume, as the prime determinant of the nanopore ionic current. At a qualitative level, the simulations reproduced the smaller blockades (larger currents) experimentally observed for homopolymers consisting of larger (dA) nucleotides, in comparison to the blockades produced by poly(dC) or poly(dT). Quantitative comparison was not possible partially because of the large statistical error in the simulated currents and, as we show below, because of force field artifacts.

Here, we reevaluate the ability of the all-atom MD method to quantitatively reproduce ionic current blockades produced by the translocation of ssDNA through a biological nanopore. We begin our examination with considering open pore conductance of MspA variants and their ion selectivity. Next, we compare the performance of the following three MD force fields— CHARMM36,^35,36^ AMBER Parmbsc1^37^ and DES-AMBER^38^—in their ability to reproduce blockade currents produced by the presence of DNA homopolymers. We then identify the most critical deficiencies in the force field that shows the closest agreement with experiment and attempt to improve that force field by means of non-bounded (NBFIX) corrections.^39^ We validate the updated force field through comparison to a new set of experimental measurements.

## MATERIALS AND METHODS

### MD Simulations

#### MD simulation of open-pore MspA systems

The simulation systems were adapted from previously described equilibrated models.^34^ Initial coordinates of MspA were obtained from the Protein Data Bank (PDB ID: 1UUN).^40^ Eight arginines at position 96 were reverted to alanine, and mutations D90N, D91N, D93N, D118R D134R and D139R, were introduced to generate the M2-NNN MspA. The full-length MspA was embedded in a 1,2-diphytanoyl-*sn*-glycerol-3-phosphocholine (dPhPC) bilayer either 110×110 Å^2^, 111×111 Å^2^, or 160× 160 Å^2^ in size. A truncated model of the nanopore was constructed by removing the vestibule region of the channel containing residues 1–84 and 120–184. This model was embedded in a 65 × 65 Å^2^ dPhPC lipid bilayer. All systems were solvated in KCl electrolyte, with the bulk concentration tuned to 1 M.

After 2,400 steps of energy minimization,^41^ each system was equilibrated for 50 ns in the constant number of particle, pressure and temperature (NPT) ensemble using the NAMD3 simulation engine.^42^ Pressure was maintained using the Nośe-Hoover Langevin piston^43^ with a period and decay of 200 and 100 fs, respectively. Multiple time stepping^44^ was used: local interactions were computed every 2 fs whereas long-range interactions were computed every 6 fs. All short-range nonbonded interactions were cut off by 1.2 nm. Long-range electrostatic interactions were evaluated using the particle-mesh Ewald method^45^ computed over a 0.1 nm-spaced grid. The 1–4 scaling factor for the electrostatic interactions was set to 1.0. SETTLE^46^ and RATTLE82^47^ algorithms were applied to constrain covalent bonds to hydrogen in water and in non-water molecules, respectively. All simulations employed periodic boundary conditions.

Production simulations were then carried out for hundreds of nanoseconds under the constant number of particles, volume and temperature (NVT) ensemble. External electric *E* was applied to produce the desired drop of voltage across the membrane as *V* = −*EL_z_*, where *L_z_* is the system’s dimension along the electric field vector. ^29^ Throughout both equilibration and production runs, Langevin thermostat^48^ with a dumping coefficient of 1 ps*^−^*^1^ was applied to all non-hydrogen atoms of the lipid membrane to maintain temperature at 295 K. All C*α* atoms of the pore were restrained to their crystallographic coordinates using harmonic potentials of 1 kcal mol*^−1^* Å*^−2^* spring constant, unless specified otherwise. The simulations used the CHARMM36 force field,^35,36^ the TIP3P water of model,^49^ the ion parameters distributed with CHARMM^50^ modified using the CUFIX corrections.^51–53^ Atomic coordinates were recorded every 9.6 ps, unless specified otherwise. Visualization and analysis were performed using VMD^54^ and MDanalysis.^55^

The study of the open pore current dependence on the system size, the transmembrane bias was 180 mV; these runs are summarized in SI Table 1. MD simulations of MspA mutants used 110×110× 120 Å^3^ (SI Table 2) and 65×65× 80 Å^3^ systems (SI Table 3) under ±200 mV bias, with all other settings identical to those employed for M2-NNN. The simulations that tested the effect of halide NBFIX^56^ (SI Table 4) used the same settings as the above. Table 1 lists the value of K^+^ and Cl*^−^* NBFIX corrections used in this work. All simulated current values listed in the tables are raw MD currents without any scaling.

**Table 1:**
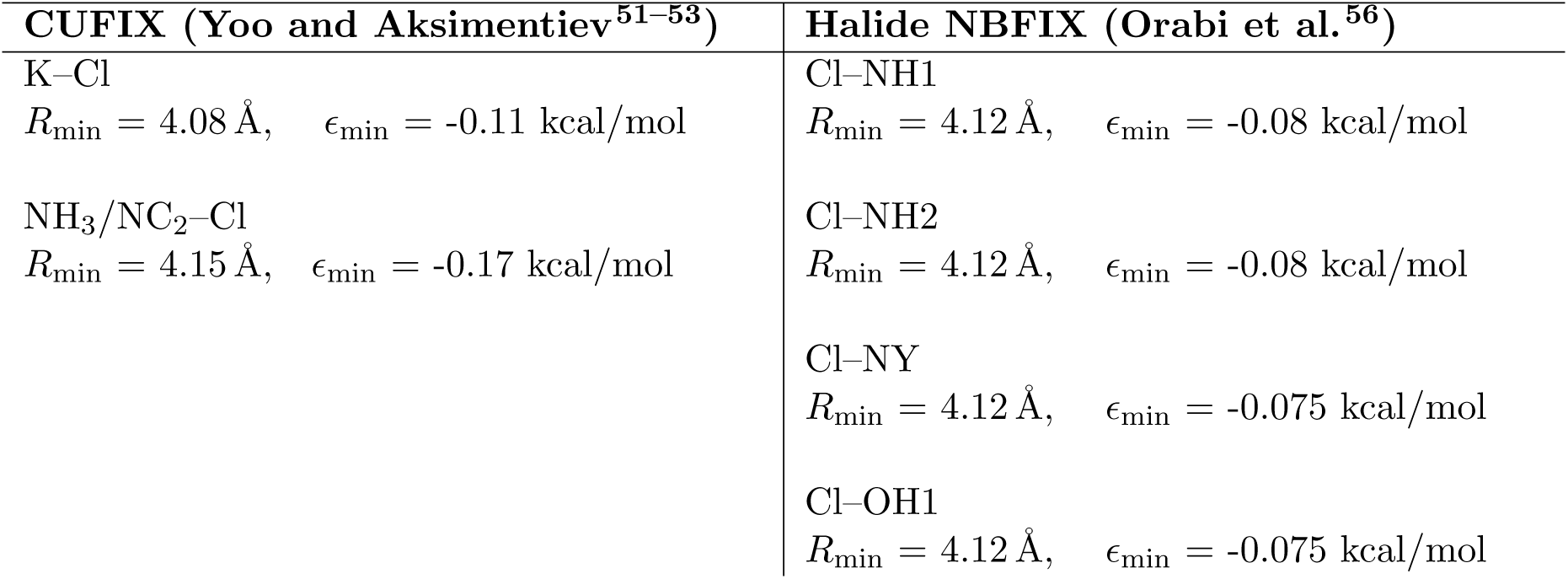
Pair-wise NBFIX corrections to ion parameters tested in this work.

#### MD simulations of ion selectivity

Under symmetric ion condition, M2-NNN MspA was simulated using severa; sets of NBFIX corrections, listed in Table 1. The simulation system was 111 × 111 × 233 Å^3^ and the simulations were conducted under a 180 mV transmembrane bias using the protocols described above.

To generate a stable ion concentration gradient across the simulation system, we employed a custom boundary constraint^57^ implemented using the tclBC interface of NAMD2.^42^ The simulation system was 180 Å in length along the *z* axis and 111×111 Å^2^ along the *x*–*y* plane. A boundary plane, running parallel to the membrane, was defined at *z*_boundary_. When an ion’s *z* coordinate exceeded that of the boundary plane, *i.e.*, *z_i_ > z*_boundary_, a constant restoring force of −0.024 kcal mol*^−1^* Å*^−1^* was applied along the *z* axis to push the ion toward the membrane. The action of the force locally increased ion concentration in the region between the boundary plane and the membrane. The ions were allowed to diffuse through the nanopore unimpeded. By tuning the magnitude of the restoring force and the location of the boundary plane, a desired gradient of ion concentration was realized.

For efficiency, the boundary force script targeted only those ions that, in the previous evaluation cycle, resided within a margin zone defined as *z_i_* ≥ *z*_boundary_ − *δz*_margin_. The use of the margin zone ensures that only the ions that are likely to cross the boundary are considered within the tclBC script, which improves the computational efficiency of the simulation. The list of ions subject to the boundary force was updated every 100 integration steps. The margin zone width was 25 Å. The total applied force was monitored and recorded every 100 steps. SI Table 5 lists all ion selectivity simulations that were performed.

#### MD simulations of poly(dT) and poly(dC) homopolymers in M1-NNN MspA

The ssDNA systems were adapted from our previous work, ^34^ extending the strands to 13 nucleotides. The bulk ion concentration was set to 1 M. The C1*^′^* atom of the 3*^′^* terminal was restrained 30 Å above the center of the MspA constriction a using a harmonic potential of 1 kcal mol*^−1^* Å*^−2^* spring constant. After a 45 ns equilibration using NAMD3, production simulations were performed under 180 mV bias on D. E. Shaw Research Anton 2 supercomputer^58^ using the CHARMM36 force field with CUFIX corrections.^53^ Long-range electrostatics were evaluated using the *u*-series Gaussian expansion.^59^ Temperature and pressure were controlled using a Nośe–Hoover thermostat and Martyna–Tobias–Klein barostat within the Multigrator framework.^60^ Unless noted otherwise, remaining parameters matched those used in the NAMD3 runs.

To determine how the blockade currents depend on a force field model, additional simulations were carried on Anton 2 using either the bsc1^37^ or DES-AMBER 3.20^38^ force fields. For bsc1 simulations, we followed the protocol from Cheatham’s force field assessment,^61^ combining ff19SB^62^ for protein, bsc1^37^ for DNA, OPC for water,^63^ and Joung–Cheatham ion parameters.^64^ For DES-AMBER simulations, we used the DES-AMBER force field^38^ for protein, DNA, and ions and the TIP4P-D water model.^65^ These simulations we conducted under the same ion concentration and applied voltage conditions as in the corresponding CHARMM runs, and employed the same thermostat, barostat, and *u*-series treatment of long-range electrostatics.

Open-pore simulations of the M1–NNN mutant using both CHARMM and AMBER force fields were performed using a simulation unit cell of the truncated DNA–MspA systems, 65 × 65 × 80 Å^3^. All simulation parameters were identical to those used in the simulations of MspA containing ssDNA strands. The resulting open-pore currents under 180 mV were 610 pA for CHARMM (TIP3P water model), 250 pA for AMBER bsc1 (OPC water model) and 390 pA for DES-AMBER (TIP4P-D). After correcting for the differences in the bulk electrolyte conductivity SI Fig. 1, the scaled open-pore M1–NNN current obtained using the CHARMM force field was 372 pA. Similarly, using the reported conductivities of 1M KCl solution simulated using the OPC water model and Joung–Cheatham ion parameters,^66^ *i.e.,*∼ 7.5 S/m, the scaled open-pore current for AMBER bsc1 was about 350 pA. The bulk conductivity of KCl within the DES-AMBER model was determined through a series of MD simulations varying the bulk electrolyte concentration and appleid electric field, SI Fig. 2. These simulation were performed using the TIP4P-D model of water^65^ and Joung–Cheatham ion parameters^64^ without any NBFIX corrections. In general, bulk conductivity of KCl within the DES-AMBER model was considerably smaller than the experimental one, in contrast to values obtained using standard CHARMM. This change that could have been expected given the higher viscosity of the TIP4P-D water in comparison to TIP3P. Using the obtained bulk conductivity of 1M KCl and after scaling the raw MD current by the ratio of experimental and simulated bulk conductivities, the scaled open-pore M1–NNN current was 719 pA. SI Table 6 lists the resulting blockade current values for the performed simulations.

#### MD simulations of osmotic pressure

The osmotic pressure was computationally determined following the protocol established elswhere.^51,67^ Each osmotic pressure simulation was performed using a system separated into two compartments by virtual semipermeable membranes oriented parallel to the *x*−*y* plane. One compartment was filled with a nucleoside solution, while the other contained pure water. Each nucleoside molecule was electrically neutral, consisting only of the sugar and the nucleobase moieties. Each semipermeable membrane was modeled using a half-harmonic planar potential that acted exclusively on the solute molecules:

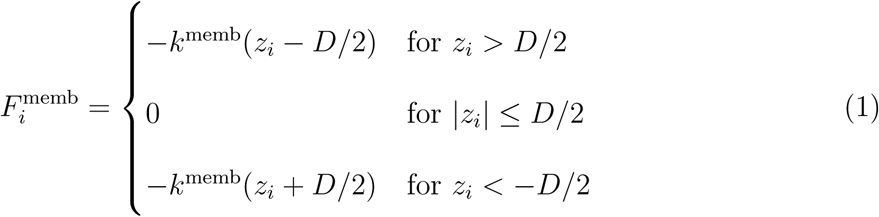

where *z_i_* is the *z* coordinate of solute *i*, *D* is the width of the nucleotide solution compartment, and the force constant *k*^memb^ = 9.56 kcal mol*^−1^* Å*^−2^*. These half-harmonic potentials were implemented in NAMD2 via the tclBC interface and applied to C1*^′^* atoms of the sugar ring of each nucleotide. For each nucleotide concentration, the system was first equilibrated for 5 ns, which was followed by three replica production runs of approximately 30 ns. During the production simulations, the instantaneous forces exerted by the membrane potentials on the nucleotides were recorded. The instantaneous pressure exerted by each membrane was computed by dividing the negative of the total force applied by the membrane to the solutes by the system’s *x* − *y*cross section area. The average osmotic pressure was determined by averaging the instantaneous pressure values over the final 50 ns of each 70 ns production trajectory and over the three replica simulations.

#### Validation simulations of asparagine–DNA backbone NBFIX

The NBFIX was validated by simulations a poly(dA)_11_ strand threaded through M1-NNN MspA in the 3*^′^*-*trans* orientation and the abasic construct dT_4_(idSp)_4_dT_4_ threaded also 3*^′^*-*trans* through M2–NNN and M2–SNN MspA. Starting from a poly(dT)_12_ conformation, extended from the poly(dT)_11_ system obtained from our previous simulations,^34^ the four nucleotides closest to the MspA constriction were converted into abasic (idSp) residues. The abasic residues for CHARMM simulations were constructed using the DELB patch, whereas dZETA parameters were used for DES-AMBER simulations. All simulations used the truncated (65 × 65 × 80 Å^3^) MspA model and followed the same protocols as our other simulations of ssDNA homopolymers. SI Table 7 provides a summary of the simulations.

#### MD simulations of NeutrAvidin-tethered ssDNA in full-length MspA

A poly(dT)_21_ strand was covalently linked to biotin through a six-carbon alkyl spacer attached to its 5*^′^*end. CHARMM-compatible topology and parameters for the biotin and the alkyl linker were generated using CGenFF.^68,69^ The biotin moiety of the construct was placed into the binding pocket of NeutrAvidin, forming the NeutrAvidin-biotin-DNA complex. This construct was then threaded through an equilibrated full-length M2-NNN MspA nanopore that was embedded in a dPhPC lipid bilayer. The entire system was re-equilibrated for 50 ns before production simulations. The production runs were performed or 2 µs on Anton 3 under 180 mV bias. A harmonic restraint (*k*_spring_ = 2.0 kcal mol*^−1^* Å*^−2^*) was applied between geometry center of biotinyl group and the NeutrAvidin Trp_97_ side chain to prevent dissociation of the biotin anchor; the equilibrium distance of the restrain was set to 6.5 Å.

#### Ionic current and electrostatics calculations

Instantaneous ionic current was calculated as^70^

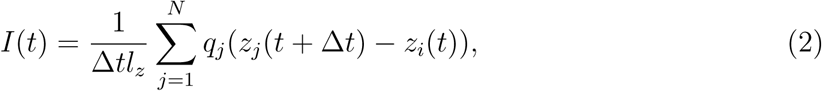

where *z_j_*(*t* + Δ*t*) − *z_i_*(*t*) is the displacement of ion *j* along the *z* direction during the time interval Δ*t* = 20 ps and *q*_j_ is the charge of ion *j*. To minimize the effect of thermal noise, the current was calculated within an *l_z_* = 20 Å thickness slab centered at the nanopore constriction; the slab spanned the entire simulation system in the *x*-*y* plane.

The 3D electrostatic potential was computed using the PMEPot plugin, ^29^ and the one-dimensional profile along the *z* axis was obtained by averaging within a central cylindrical region of 4 Å radius.

#### Radius profile of the pore

At each position along the *z* axis within the pore region, a “test” radius was initialized at zero and gradually increased until the corresponding cross-sectional circle intersected with the pore wall defined as the point where non-hydrogen atoms of the protein first appear. The radius at which that intersection occurred was recorded as the pore radius at that specific *z* coordinate.

#### Base stacking

Base stacking was quantified using cpptraj.^71^ For each pair of neighboring bases, we computed the inter-base distance and the angle between their plane normals. Two bases were considered stacked if their inter-base distance was less than 5 Å and the angle between their plane normals was below 45*^◦^*. Only adjacent bases along the strand were included in the analysis.

#### SHapley Additive exPlanations Analysis

Frames were labeled as “good” or “bad” based on the agreement between the SEM blockage current^72^ and the experiment blockage current. To enhance the contrast between the “good” and “bad” frames, only the frames that fall within close range of the experimental current and those ones significantly deviating from experimental values were selected. In the analysis for DNA homopolymers, CatBoost classification model^73^ was applied to thousands of conformations. In the SHAP method, features were bootstrapped and those strongly correlated with high-scored predictions were ranked high in the SHAP evaluation. The analysis was conduced using the SHAP package for python.^74^

### Experiment

#### Materials

All chemicals were purchased from Thermo Fisher Scientific unless otherwise specified. LB-medium and LB agar were purchased from Boston BioProducts. Ampicillin and acrylamide/bis-acrylamide 29:1 solution were purchased from Research Products International. BL21 (DE3) pLysS *E. coli* cells were purchased from Invitrogen. dPhPC lipid was purchased from Avanti Polar Lipids. Streptavidin was purchased from Bio Basic. Plasmid purification miniprep kits were purchased from New England Biolabs. Mini-PROTEAN TGX Stain-Free gels were purchased from BioRad. Unstained broad-range protein standard and DpnI restriction enzyme were purchased from New England BioLabs. Phenylmethylsul-fonyl fluoride (PMSF) was purchased from Research Products International. All oligos tested were purchased from Integrated DNA Technologies (IDT). DNA plasmid encoding M2-NNN MspA was a gift from Dr. Giovanni Maglia (University of Groningen, Netherlands).

#### Mutagenesis

Mutations were introduced into the MspA gene by overlapping mutagenesis PCR using a pT7 plasmid containing the M2-NNN MspA as a template. The PCR products underwent a DpnI digestion for 3 h at 37 *^◦^*C to remove the template DNA plasmids. After 20 min incubation at 80 *^◦^*C to deactivate the DpnI enzyme, the reaction mixture was transformed into chemically competent DH10*β E. coli*. Colonies containing the desired mutation were verified by sequencing (Eurofins Genomics). Primers were purchased from Eurofins.

#### Purification of mutant MspA nanopores

Plasmids containing pT7-M2 MspA mutants were transformed into BL21 (DE3) pLysS cells and grown in LB medium at 37 *^◦^*C until OD_600_ reached 0.5–0.6. The cells were induced with 0.5 mM IPTG and continued to grow at 16 *^◦^*C for ∼16 h. Cells were harvested by centrifugation at 4000 rpm for 15 min at 4 *^◦^*C. Cells were resuspended in lysis buffer (100 mM Na_2_HPO_4_/NaH_2_PO_4_, 1 mM EDTA, 150 mM NaCl, 1 mM PMSF, pH 6.5) and heated at 60 *^◦^*C for 10 min. Cells were placed on ice and sonicated using a VWR Scientific Branson 450 sonicator (duty cycle of 20% and output control of 2) for 10 min. The lysate was centrifuged at 13,000 rpm for 20 min at 4 *^◦^*C and the supernatant was discarded.

The pellet was resuspended in solubilization buffer (100 mM Na_2_HPO_4_/NaH_2_PO_4_, 1 mM EDTA, 150 mM NaCl, 0.5% (v/v) Genapol X–80, pH 6.5). After complete resuspension, the sample was mixed for∼ 30 min and centrifuged again at 13,000 rpm for 20 min at 4 *^◦^*C. The supernatant containing the solubilized membrane extract was purified using a gravity Ni-NTA affinity column. The column-bound protein was washed with buffer A (0.5 M NaCl, 20 mM HEPES, 0.5% (v/v) Genapol X–80, pH 8.0), buffer A1 (same as buffer A plus 50 mM imidazole), and eluted with buffer A2 (same as buffer A plus 200 mM imidazole).

The eluted MspA was run on a 7.5% SDS-PAGE gel. The band corresponding to the MspA octamer was cut from the gel, crushed, and incubated with extraction buffer (50 mM Tris-HCl, 150 mM NaCl, 0.5% Genapol X–80, pH 7.5) at room temperature for 2 h. After centrifugation at 14,000 rpm for 25 min at 4 *^◦^*C, the supernatant containing the extracted MspA oligomer was confirmed by a 7.5% SDS-PAGE gel. Proteins were frozen in liquid nitrogen and stored at −80 *^◦^*C until use.

#### Single-channel current recordings

Recordings were performed at room temperature (22 *^◦^*C). Experiments were carried out in a custom chip consisting of two chambers separated by a 25 *µ*m-thick Teflon™ film with a central aperture of ∼ 70 *µ*m in diameter. The aperture was pretreated on both sides with 10% (v/v) hexadecane in pentane. Both chambers were filled with 900 *µ*L buffer (10 mM HEPES, 1 M KCl, pH 8.0). dPhPC dissolved in pentane (10 mg mL^−1^) was added to the surface of the buffer in both chambers. Ag/AgCl electrodes were immersed in each chamber, with the cis side grounded, and an artificial lipid bilayer was formed by pipetting the solution in both chambers below the aperture several times. The M2-MspA mutant protein was added to the cis chamber. Upon single-pore insertion, 100 nM streptavidin and 300 nM 5*^′^*-biotinylated ssDNA were added to the cis chamber. Each ssDNA sample was subjected to repeated cycles of a sweep-voltage protocol for 10 min, with each cycle consisting of a 10 s hold at +150 mV to collect signature current levels followed by a 200 ms pulse at −150 mV to release the ssDNA from the pore. Current signals were amplified with an Axopatch 200B patch clamp (Axon Instruments), filtered with a 2 kHz Bessel filter, and acquired by a Digidata 1440A board (Axon Instruments) at a sampling rate of 50 kHz. Data analysis was performed using Clampfit 11.2 (Molecular Devices).

#### Ion selectivity

The reversal potentials of the MspA variants were determined as follows: the grounded cis chamber was filled with 900 *µ*L of 3 M KCl and 10 mM HEPES (pH 8.0), and the trans chamber was filled with 900 *µ*L of 1 M KCl and 10 mM HEPES (pH 8.0). Ag/AgCl electrodes were connected to each chamber via a 0.4% agarose salt bridge prepared in 3 M KCl. MspA proteins were added to either the cis or the trans chamber. Upon single-pore insertion, a reversal potential was applied to set the current to zero. Ion selectivity was calculated from the measured reversal potential using the Goldman–Hodgkin–Katz (GHK) equation.

## RESULTS

### Open pore current of MspA nanopore

We begin our inquiry by re-examining the open pore conductance of MspA mutants. The wild-type (WT) MspA structure contains three rings of aspartic acid side chains at position 90, 91 and 93, which places 24 negative charges in the narrowest portion of its transmembrane pore.^11^ Not surprisingly, WT MspA exhibits pronounced gating which makes it ill-suited for nanopore sensing applications. However, replacing the three rings of aspartic acids with asparagines virtually eliminates the gating behavior. Thus, several variants of such D90N/D91N/D93N mutants—known as NNN MspA pores—have been developed, differing by the placement of additional arginine residues in the MspA vestibule to facilitate ssDNA capture.^75^

In our previous MD study,^34^ the simulated open-pore current through full-length NNN mutant of MspA (the M1-MspA nanopore) was 610 pA at 180 mV bias and 1M KCl bulk electrolyte. Even greater open pore current (910 pA) was recorded from a simulation of a minimal (truncated) model of MspA that contained only the transmembrane part of the nanopore. ^34^ Experimentally, open-pore current through full-length M1-NNN nanopore under identical bias and electrolyte conditions was measured at 328 pA. ^21^ At first look, the difference between simulation and experiment is substantial and greatly exceeds the simulation-versus-experiment differences seen for other nanopores: alpha-hemolysin,^29^ OmpF,^76,77^ OmpC^78^ and aerolysin.^79^ One should, however, recall that the all-atom MD method systematically overestimates bulk conductivity of dense electrolyte solutions ^29,80^ by a factor that depends on the specifics of the water model, thermostat, the cutoff scheme for evaluation of vdW and electrostatic interactions, etc.^81^ A practical solution is to scale the simulated nanopore current *I*_raw_ by the ratio of experimental (*σ*_exp_) and simulated (*σ*_sim_) bulk electrolyte conductivity: *I*_corr_ = *I*_raw_ × *σ*_exp_*/σ*_sim_. However, even after taking such a correction, the difference (∼ 24%) remains substantial. In our previous study,^34^ we attributed the remaining difference to access resistance that is present in experiment but is minimal in an MD simulation.^82^ Below, we re-examine the ability of brute-force MD to reproduce the experimental current through open MspA nanopore.

For these studies, a full-length M2 MspA nanopore—the NNN mutant containing an additional ring of arginine side chains at position 118 above the nanopore constriction^75^—was embedded in a lipid membrane and immersed in 1M KCl solution, Fig. 1a. To test the effect of access resistance, three systems containing full-length M2 MspA were constructed: the smallest system that could accommodate a full-length MspA nanopore, 110 × 110 × 120 Å^3^ (System I, 152,763 atoms), a larger and higher aspect ratio system, 160 × 160 × 200 Å^3^ (System II, 546,222 atoms), and a high aspect ratio system, 111 × 111 × 291,923 atoms). The fourth system contained a truncated model of M2 MspA embedded in a 65 × 65 × 80 Å^3^ unit cell (System IV, 37,259 atoms). For these simulations, the protein nanopore and the lipid (dPhPC) membrane were described using the standard CHARMM36 force field^35,36^ in tandem with a TIP3P water model^49^ which rigid bonds were enforced using the SETTLE algorithm.^83^ The ion parameters distributed with CHARMM^50^ were modified using the CUFIX corrections ^53^ that reproduce the osmotic pressure of KCl and potassium acetate solutions,^51^ with the latter being a model for K^+^ interaction with the aspartic and glutamic side chains of MspA. Test 20 ns simulations of a 23.9 × 23.9 × 23.9 Å^3^ electrolyte volume under 200 mV and 400 mV electric bias determined the simulated bulk conductivity of 1 M KCl electrolyte as 17.2 S/m at 295 K, SI Fig. 1. For reference, the experimental bulk conductivity of KCl electrolyte^80^ at 295 K is 10.5 S/m. Please note that the application of CUFIX corrections for K^+^–Cl*^−^* interactions^51^ increases the bulk conductivity value slightly from that reported in Ref. 81

**Figure 1:**
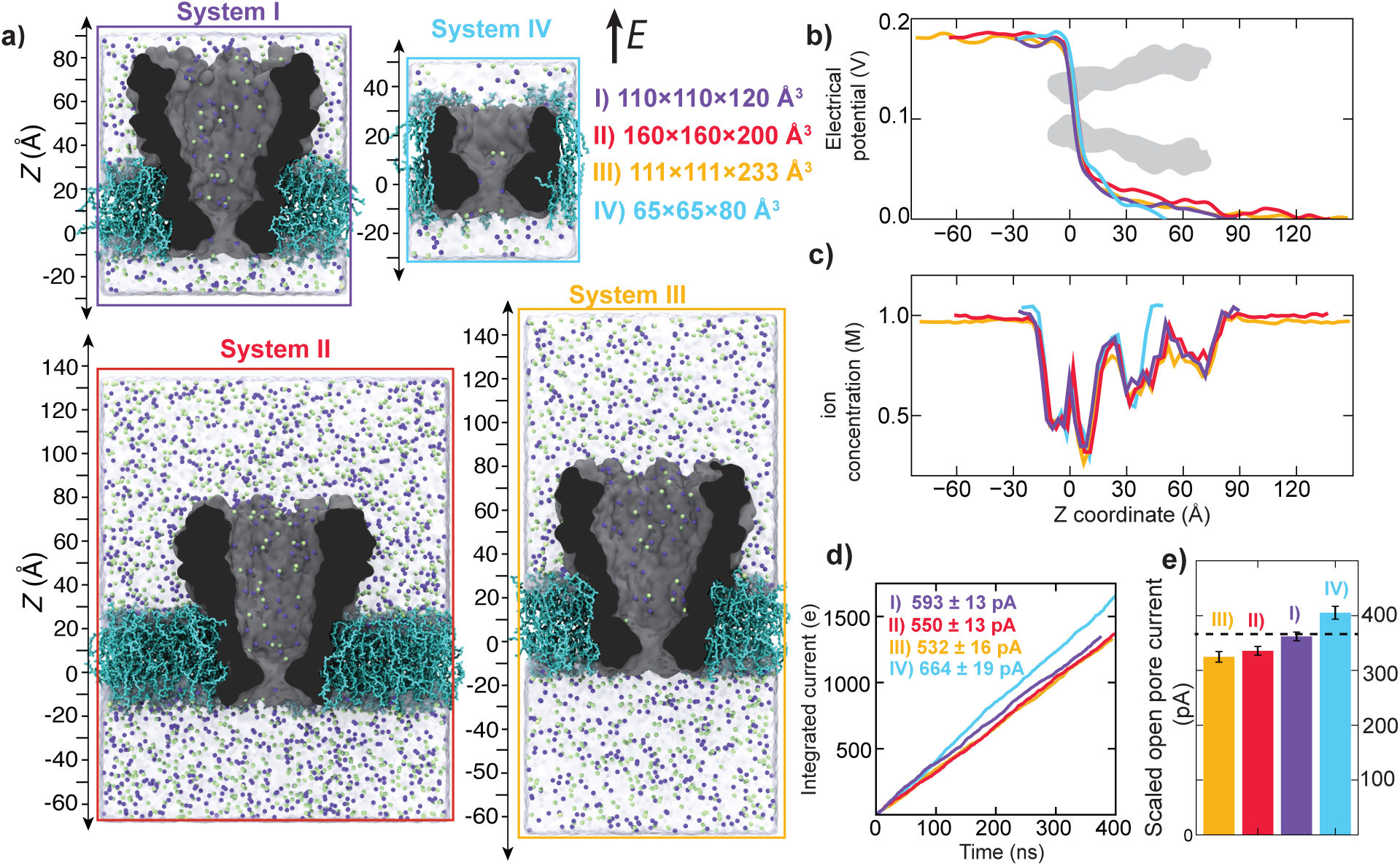
Open-pore simulations of M2-MspA nanopore. **a,** Cut-away view of all-atom M2-MspA models each containing one nanopore (gray), a lipid bilayer membrane (cyan), explicit water (semi-transparent molecular surface), potassium (purple) and chloride (green) ions. The systems’ dimensions color-match the corresponding data in subsequent panels. **b,c** Average electrostatic potential (panel b) and ion concentration (panel c) along the nanopore axis, *Z*, defined in panel a. The electrostatic potential was computed by averaging over the corresponding simulation trajectory (sampled every 50 ps) within 4 Å radius cylindrical bins stacked in 1 Å intervals along the pore axis. The inset image of the MspA pore aligns with the *Z* axis. Ion concentration profiles were averaged using 3 Å spaced bins over 32 Å radius cylinders for *Z* values laying within the nanopore volume and over the entire extent of the system outside the nanopore. **d,** Total electrical charge transported by ions through the nanopore *versus* simulation time. The slope of each line is the average current. Mean and standard error values were computed by splitting MD trajectories into 50 ns intervals. **e,** Mean simulated open-pore current after scaling the raw MD current by 0.61, the ratio of experimental and simulated bulk conductivity of 1 M KCl at 295 K. The dashed line indicates the open pore conductance of M2 NNN MspA measured in this work, 370 pA.

Upon minimization and equilibration, the four systems were simulated under external electric field applied normal to the lipid membrane, choosing the strength of the field in each system to produce the same 180 mV transmembrane voltage. Indeed, the average distribution of the electric potential along the nanopore axis, Fig. 1b, exhibits a 180 mV drop across all four systems. The average profile of the electrostatic potential along the nanopore axis has the same shape in all four systems, with the sharpest drop of the potential occurring at the narrowest part of the nanopore. The local ion concentration exhibits reproducible variation within the nanopore volume regardless of the system size, Fig. 1c, returning to a bulk value away from the nanopore.

The application of the external field produced displacements of ions through the nanopore, which we quantify by plotting the total charge transported by the ions as a function of simulation time, Fig. 1d. The slope of the lines—the average ionic current—is seen to depend on the system size, SI Table 1. In line with results of a previous study, ^82^ the system’s aspect ratio was found to affects the simulated value of the ionic current, with the highest aspect ratio system (System III) having the highest resistance (compare System I and III data). The ionic current measured for System I matched the experimental value within the statistical error of the current determination after scaling the simulated current with the ratio of experimental and simulated bulk conductivities, Fig. 1e. Interestingly, the current for the truncated MspA system (System IV) was not too far off from the experimental one even though a significant portion of the nanopore was removed. Given their modest atom count, Systems I and IV provide excellent platforms for MD study of blockades currents.

A closer examination of the ion concentration profiles, Fig. 1c, reveals that the local ion concentration away from the nanopore does not match exactly the prescribed bulk concentration, *i.e.*, 1 M, despite all systems having exactly the same ion-to-water ratio. This effect is caused by the non-trivial variation of ion concentration within the MspA nanopore, which, given its shape, occupies a considerable fraction of the unit cell volume. Thus, if quantitative agreement with experiment is desired, one should examine and adjust the ion concentration iteratively after several test runs under external electric field. As shown in SI Fig. 3, setting the overall electrolyte concentration to 1 M using, for example, the VMD’s Autoionize plugin,^54^ can result in bulk concentrations much greater than 1 M, which leads to a considerable overestimation of the simulated current. Scaling the MD current with the experiment-to-simulation ratio of bulk conductivities using the actual, observed concentration of ions in the bulk regions of the simulated system can largely correct the overshot, SI Fig. 3.

### Open pore current of MspA mutants

To test the robustness of the MD approach in describing the ion conductance of engineered nanopores, we experimentally expressed a library of mutant M2 MspA and measured their open pore conductance. Specifically, one or two rings of asparagine (N) residues at the nanopore constriction, Fig. 2a, were mutated into either glycine (G) and/or serine (S) residues producing the following six M2 variants: GNS, GSN, NGS, NNS, NSN and SNN, where the three letter code indicate the side chains located at positions 90, 91 and 93, respectively. Three additional mutants were prepared varying the side chain of residues 91: NLN, NAN and NGN. Following the standard electrophysiology procedures, the open pore current was measured under ±200 mV, see Methods for details. Figure 2b shows typical current traces recorded for the GNS and SNN mutants. Despite frequent gating, the open pore conductance could be accurately determined from the all-point histogram of the current. Repeating the measurements for all ten M2 variants revealed a pronounced dependence of the current on the specific mutation, Fig. 2c.

**Figure 2:**
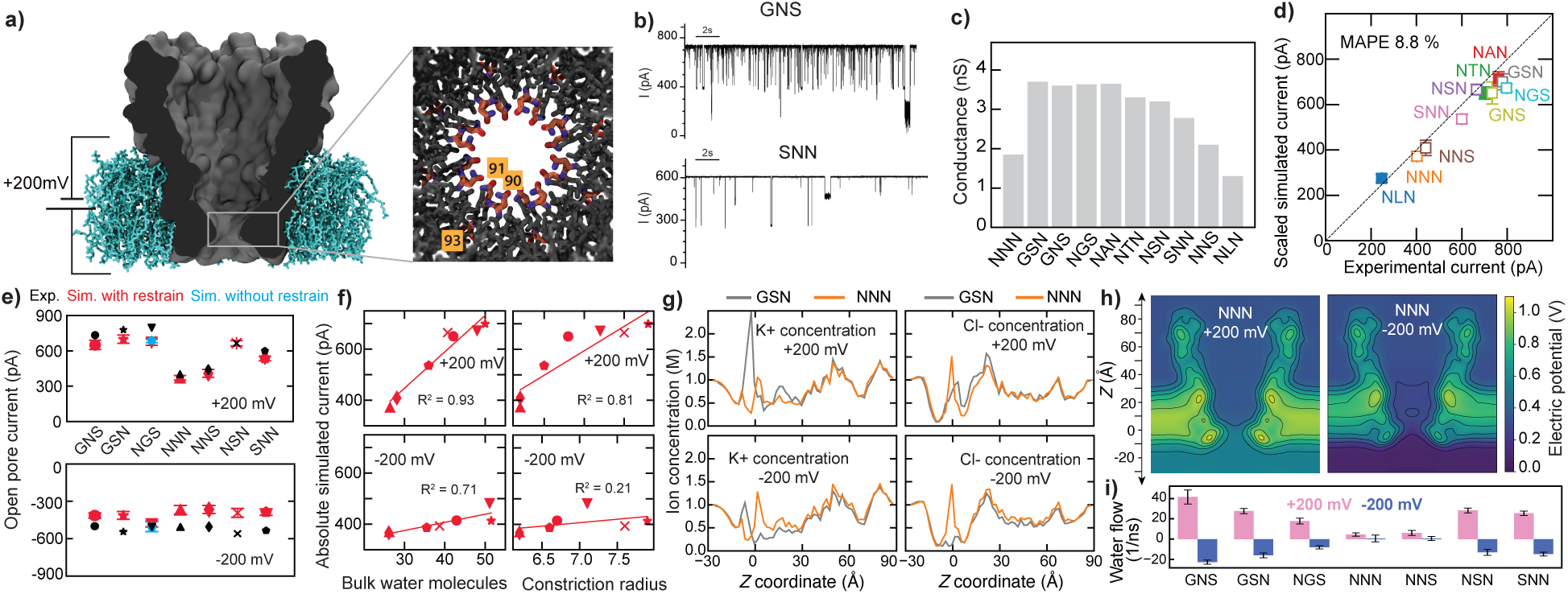
MD simulations MspA mutants. **a,** All-atom model of M2 MspA. The inset shows a top-down view of the pore with the mutation sites 90, 91, and 93 highlighted. **b,** Representative ionic current recording from M2 90G/91N/93S (GNS) and M2 90S/91N/93N (SNN) mutants experimentally measured at 10 mM HEPES, pH 8.0, 1.0 M KCl and at +200 mV. **c,** Experimental conductance of MspA mutants. **d,** Simulated *versus* experimental open-pore currents at +200 mV. Open and filled symbols indicate MD data obtained using 270 ns simulations of System I and 300 ns simulations of System IV, respectively. Error bars show standard deviations calculated by splitting each trajectory into 50 blocks and treating the average within each block as an independent measurement. Mean absolute percentage error (MAPE) of scaled MD current with respect to experiment is 8.8%. **e,** Open-pore current of seven MspA variants measured in experiment (black) and simulation (red/blue) under ±200 mV. In all simulations, C*α* atoms of MspA were restrained to their crystallographic coordinates except for the two simulations (blue) that were carried out in the absence of any restrains. **f,** Magnitude of the simulated current *versus* number of bulk-like waters in the MspA constriction (left) and *versus* its radius (right). Red lines indicate linear fits to the data. Symbols are defined in panel e. Top / bottom row show data at +/−200 mV. **g,** Trajectory-averaged concentration of K^+^ and Cl*^−^* ions along the axis of the GSN and NNN pore. **h,** Electrostatic potential in M2 NNN MspA under ±200 mV. The maps, averaged over the MD trajectory and the cylindrical symmetry of the nanopore, are shown for a plane encompassing the nanopore axis. **i,** Average rate of water flow through mutant pores at ±200 mV. Each value is a 270 ns trajectory average and the error bars denote the standard error of mean computed using 20 ns blocks.

Starting from a well-equilibrated configuration of System I containing the NNN variant of MspA, Fig. 1a, we constructed six additional systems by mutating the side chains of the equilibrated structure according to the sequence of the experimentally tested mutants. After 50 ns equilibration, each system was simulated under a 200 mV transmembrane bias of either polarity for 270 ns using the CHARMM36 force field and the CUFIX corrections to K^+^ parameters. Three additional mutants (NLN, NAN and NGN) were prepared using the truncated MspA model (System IV) and simulated for 300 ns under +200 mV using the CHARMM36 force field and CUFIX corrections. Upon appropriate scaling of the raw MD current with the ratio of experimental and simulated bulk conductivities, the absolute value of the simulated current appear to closely follow that measured in experiment, Fig. 2d, with the mean absolute percentage error of the simulated *versus* experimental current of only 8.8%. This result is reassuring as it shows the MD method has the sensitivity required to capture the effect of such minute structural changes.

Experimental data at +200 mV reveal a factor 2 change of the ionic current through the MspA nanopore upon seemingly minor changes of its structure, Fig. 2e (top) and SI Table 2. Similar variation of the open pore current were observed in the corresponding MD simulations, Fig. 2e. As previously suggested,^34^ the simulated current shows strong correlation with the number of bulk-like water molecules within the MspA constriction, its conductive volume, Fig. 2f (left), suggesting that the current differences are primarily influenced by the pore size, Fig. 2f (right).

Much smaller variations of the current were observed at a negative 200 mV bias, Fig. 2e (bottom) and SI Table 2. As a consequence, some of the mutants—GNS, GSN, NGS and NSN—manifested pronounced rectification of the ionic current, in comparison to a mildly rectifying NNN nanopore. Bulk water analysis shows a much weaker correlation with the current Fig. 2f, suggesting that effects other than nanopore geometry are at play. Indeed, close examination of select K^+^ and Cl*^−^* concentration profiles reveal a pronounced variation of the local concentration among the nanopore axis and an even stronger dependence on the polarity of the bias, Fig. 2g. At +200 mV, the GSN pore retained more K^+^ ions than the NNN pore upstream from the constriction, at −10 *< Z <* 0 Å, Fig. 2g (top left) whereas the NNN pore retained more Cl*^−^* ions in the same region under a reversed bias (−200 mV), Fig. 2g (bottom right). The 2D maps of the local electrostatic potential of the NNN pore at both voltage polarities, Fig. 2h, revealed the expected reversal of the sharp potential gradient at the nanopore constriction, although the two maps differ substantially to be mirror images of one another. To complicate the matter further, both the bias polarity and the point mutations were found to modulate the water flux through the nanopore, *i.e.*, the electro-osmotic effect, Fig. 2i, with modest changes in nanopore geometry leading to a 5-to-10 fold increase of the water flow. While deciphering the intricate relationships between the molecular structure of the nanopore, its ion selectivity and the electro-osmotic effect is beyond the scope of this work, we note that properly calibrated all-atom MD simulations can provide high-quality data for building predictive theoretical models of nanopore transport.^84^ Subject to stochastic forces, biological nanopores undergo conformational fluctuation about their average structure, the structure typically captured by X-ray crystallography or high-resolution cryogenic microscopy. At any given time, the cross section of a biological nanopore is not perfectly circular and often has a shape of an ellipse that can shift its alignment among the equivalent nanopore monomers at the time scale of nanoseconds. ^29^

Furthermore, our lab previously observed sporadic and irreversible dislocation of nanopore loops out of their equilibrium structure in *α*-hemolysin and MspA on the time scale of several microseconds. Restraining the C*α* atoms of the nanopore to their equilibrium coordinates (typically taken from experiment) has become a practical solution to the need of sampling such structural transitions, which would require much longer simulations. To assess the impact of such restrains, we simulated the ionic current through the NGS mutant at +200 mV. The open-pore currents obtained with and without the C*α* restrains were indistinguishable within the statistical accuracy of our simulations, Fig. 2e. We arrived with the same conclusion after analysis of additional simulations reported in SI Fig. 4a. Thus, restraining C*α* atoms to their equilibrium coordinates is an acceptable approximation for MD simulations of nanopore current.

Finally, we evaluated a recent force-field refinement^56^ designed to correct the underestimation of chloride–protein interactions in the standard CHARMM force field. A set of simulations was carried out to compute the ionic current through seven MspA variants at ±200 mV using the CHARMM36m force field supplemented by both the CUFIX corrections to K^+^ interactions with the nanopore and Cl*^−^* ions,^51^ and the Orabi et al.^56^ NBFIX corrections to Cl*^−^* interactions with the amino acids of the nanopore. The addition of the Cl*^−^* corrections did not significantly altered the overall values of the nanopore currents, SI Fig. 4a,b and SI Table 4. However, the ion distributions changed substantially: Cl*^−^* displayed enhanced interactions with several asparagine residues inside the MspA constriction, SI Fig. 4c. Similar increase of Cl− affinity was observed throughout the nanopore surface, SI Fig. 4e. As we show in the next section, the increased interactions of Cl*^−^* with the nanopore surface alters the ion selectivity of the nanopore, an effect that we quantitatively assess through experimental measurement and simulation.

### Ion selectivity of MspA pores

The M2-NNN mutant of MspA was engineered to have rings of positively charged amino acids at the inner surface of the nanopore to promote DNA capture, which also made the pore anion selective, Fig. 3a. We experimentally measured the ion selectivity under a salt concentration gradient: 3M KCl was added on the *cis* side of the nanopore whereas the *trans* side contained 1M KCl. The concentration gradient produces a net flux of ions through the nanopore. For an ion selective pore, the flux generates an ionic current, which in experiment can be neutralized by means of an external transmembrane potential. The voltage at which the net current becomes zero is the reversal potential, which sign and value reveal both the direction of selectivity (cation or anion) and the permeability ratio of the ions, SI Table 5. Under these conditions, the wild-type NNN variant was weakly anion-selective. Single-residue substitutions had distinct effects on selectivity: NNS exhibited increased anion selectivity, whereas NGS showed a pronounced enhancement in cation selectivity.

**Figure 3:**
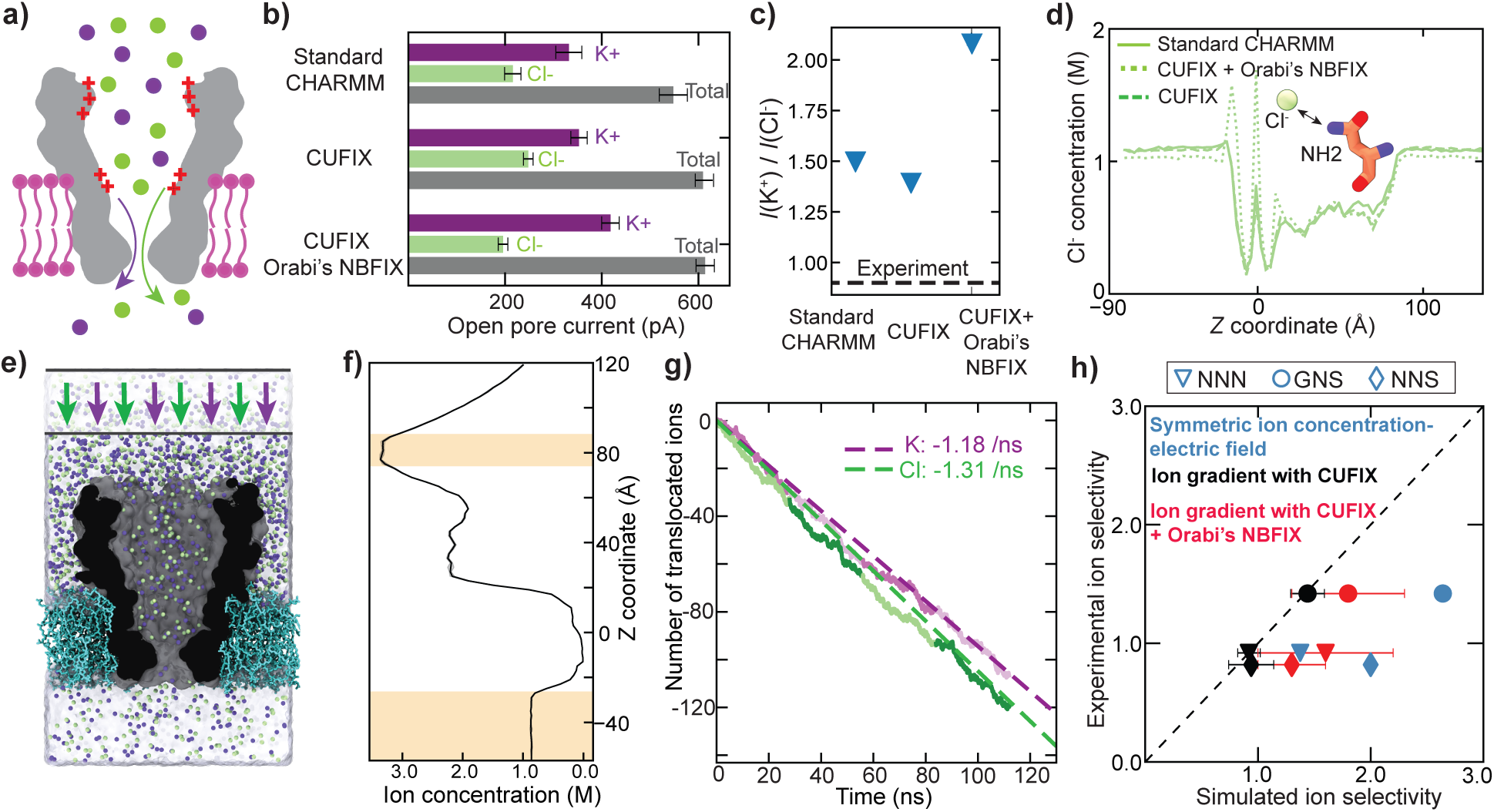
Ion selectivity of MspA mutants. **a,** Schematics of M2 MspA ion selectivity mechanism. The plus signs indicate patches of positive charge at the surface of the nanopore. Purple and green spheres depict K^+^ and Cl*^−^* ions, respectively. **b,** Average total current and the current carried by K^+^ and Cl*^−^* ions through M2-NNN MspA under +200 mV when simulated using standard CHARMM36, CHARMM36 with Yoo’s CUFIX corrections,^51^ and CHARMM36 with both CUFIX and Orabi’s NBFIX corrections. ^56^ Error bars represent the standard deviation of mean calculated using 50 ns blocks of the respective 300 ns trajectories. **c,** Simulated ion selectivity defined as the ratio of K^+^ and Cl*^−^* currents. The horizontal line indicates the experimental value obtained from reverse potential measurements performed at 3M (*cis*) /1M (*trans*) gradient of KCl. **d,** Trajectory-averaged concentration of Cl*^−^* along the pore axis. Inset illustrates the interaction of Cl*^−^* with an NH_2_ group of an asparagine in the pore constriction. **e,** Simulation setup reproducing experimental ion selectivity measurement. K^+^ and Cl*^−^* ions that cross the upper boundary (schematically illustrated) are pushed downward by an external force, creating a transmembrane gradient of ion concentration. **f,** Steady-state profile of ion concentration produced by the application of the external force. **g,** Number of K^+^ and Cl*^−^* ions translocating through the pore from the 3 M to the 1 M compartment. Data from four independent simulations (colors) are concatenated into a single trace. A dashed line represents a linear fit to a concatenated trace. **h,** Experimental *versus* simulated ion selectivity, expressed as the ration of permeability, *P* (K^+^)*/P* (Cl*^−^*). Each data point represents the average over four replica simulations. The error bars denote the standard deviation across the replicas. Experimental ion selectivity was calculated from the reverse potential measurement under 3M/1M *cis*/*trans* KCl concentrations; 10 mM HEPES, and pH 8.0. In those experiments, the MspA was added to the *cis* side.

A straightforward method of assessing the ion selectivity of a biological nanopore *in silico* is to compare the relative magnitudes of the average current carried by the respective ion species, *I*(K^+^)/*I*(Cl*^−^*).^29^ We thus analyzed our MD trajectories of the MspA mutants to compute the ratio of K^+^ and Cl*^−^* currents for the simulations carried out using the standard CHARMM36 force field,^35,36^ the CHARMM36 force field with the CUFIX corrections^51,53^ that reduce cation–protein interactions, and the CHRMM36 force field with both the CUFIX and Orabi’s NBFIX^56^ corrections that enhance anion–protein interactions. Table 1 lists the corrections used in our simulations. Under the symmetric 1 M KCl conditions and a +200 mV transmembrane bias, the M2-NNN MspA nanopore is found to be cation selective, with the selectivity ratio ranging from about 1.5 for CHARMM, 1.3 for CUFIX and 2.2 for CUFIX + Orabi’s NBFIX, Fig. 3b,c and SI Table 5. Thus, the cation selectivity was the most pronounced when using the Orabi’s NBFIX corrections. At the microscopic level, the latter NBFIX corrections caused accumulation of the Cl*^−^* ions near the nanopore constriction, particularly around the asparagine residues. This local “sticking” effect, visible as sharp peaks in the local Cl*^−^* concentration profiles, Fig. 3d, rendered the nanopore constriction negatively charged, favoring potassium transport, similar to the sticky ion action previously described for guanidinium.^85^ The accumulations of Cl*^−^* ions was not observed when using the standard CHARMM with or without the CUFIX corrections, Fig. 3d. Most importantly, the three force field models appear to produce a qualitatively inaccurate description of ion selectivity: the M2-NNN MspA was experimentally determined to be mildly Cl*^−^* selective, Fig. 3c and SI Table 5.

However, the above computational approach examines the ion selectivity of a nanopore when the ion concentration is equal on both sides of the membrane. In a way, this condition is the most relevant one for nanopore transport experiments as they typically involve symmetric buffer conditions. At the same time, it is well known that the experimentally determined selectivity ratio depends on the magnitude of the concentration gradient and can change considerably by simply changing the polarity of the concentration gradient, presumably because of the asymmetric distribution of charges among the nanopore surface. ^85^

To better reproduce the ion selectivity experiments in our MD simulations, we realized the asymmetric ion concentration conditions in a single membrane system using a method described previously.^57^ Briefly, an external force was applied to both ion types in a region near the system’s boundary to push them torward the membrane, increasing the local ion concentration on that side of the membrane, Fig. 3e. By tuning the force magnitude and the total number of ions in the system, one can achieve the desired *cis* and *trans* concentrations, which in our case were approximately 3 to 1M, Fig. 3f.

Using this refined simulation setup, we conducted four replica simulation of M2-NNN MspA and measured the ionic flux through the nanopore constriction driven by the concentration gradient, Fig. 3g. The ion selectivity simulated under these conditions showed good agreement with experiment, Fig. 3h. We then repeated this procedure for three other MspA mutants and observed much improved agreement across the board in comparison to the simulations carried out under symmetric ion conditions, Fig. 3h. Using the ion gradient protocol, we could reproduce the increased anion selectivity of the NNS mutant and the pronounced cation selectivity of the GNS mutant. Repeating ion concentration gradient simulations with the additional Orabi’s NBFIX ^56^ produced noticeable deviations of the simulated selectivity ratios from experiment, Fig. 3h, which suggest that that refinement may have overestimated the interaction between chloride and the protein. These results highlights the need for systematic benchmarking of MD force fields^66^ with respect to experiment in a manner that makes direct comparison of the results possible.

### Blockade current from ssDNA in three force field models

In our previous work, ^34^ we used the CHARMM36 force field to simulate ionic current blockade from DNA homopolymers poly(dT), poly(dC), and poly(dA) threaded through a truncated model of M1-NNN MspA in both global orientations. The simulations were carried out under the experimental voltage of 180 mV using the DE Shaw Anton machine, ^86^ which allowed us to sample the systems’ dynamics on the times scale of tens of microseconds. The simulations revealed pronounced fluctuations of the current caused predominantly by the conformational dynamics of the DNA strand. The fluctuations of the current were highly correlated with the fluctuations of the nanopore’s conductive volume—the number of bulk-like water molecules in the MspA constriction. Averaging the currents over the MD trajectories of the 5*^′^*-*trans* threaded ssDNA produced a reasonable qualitative agreement with experiment:^21^ the poly(dA) current was the highest, the poly(dT) current was the lowest and one of poly(dC) was somewhere in between.^34^ However, the average currents were clearly in quantitative disagreement with experiment, in particular for the 3*^′^*-*trans* threaded poly(dA), where a complete current blockade was observed. While we could not rule out insufficient sampling as a root cause of the discrepancy, a more likely factor was the imperfect parameterization of ssDNA, which had previously invited several computational studies.^87,88^ In this section, we evaluate the performance of three MD force fields for their ability to reproduce the experimental ionic current blockade data. We narrow the scope of our investigation to the M1-NNN MspA nanopore, which was employed in the early experimental study.^21^ First, we repeat the simulations of poly(dT) reported in our earlier work^34^ after extending the strand to thirteen nucleotides and harmonically restraining the C1*^′^* atom of its 3*^′^*-*trans* terminal to a position 30 Å above the nanopore constriction along the nanopore axis, Fig. 4a. The use of the restrain reproduces the action of a streptavidin protein that, in experiment, prevents the passage of ssDNA through the nanopore, giving ample time to obtain accurate reading of the current blockade. ^21^ The simulations of poly(dT)_13_ were run under 180 mV for 10 *µ*s using the D.E. Shaw Anton 2 machine^58^ and the CHRMM36 force field with the CUFIX corrections.^51–53^ Two additional simulations were conducted using two newer AMBER force fields: AMBER parmbsc1,^37^ a popular and extensively validated model for nucleic acids^61,89,90^ and DES-AMBER,^38^ a force field optimized to describe, among other targets, the interactions of proteins with DNA. The CHARMM simulations employed the TIP3P water model^49^ in tandem with the CHARMM-compatible Beglov–Roux ion parameters^91^ and the CUFIX corrections.^53^ The DES-AMBER simulations were conducted using the TIP4P-D water model^65^ and the corresponding DES-AMBER ion parameters, whereas the AMBER parmbsc1 simulations used the OPC water model^63^ with the Joung-Cheatham ion parameters.^64^ Additional simulation details are provided in SI Table 6 and Methods. The three MspA nanopore systems were also simulated in the absence of DNA under otherwise identical conditions, giving us the average open pore current, *I*_O_. This allowed us to report the outcome of the blockade current simulations using the relative blockade current (*I/I*_O_), eliminating the need to scale raw MD currents with the ratio of experimental and simulated bulk ion conductivities.

**Figure 4:**
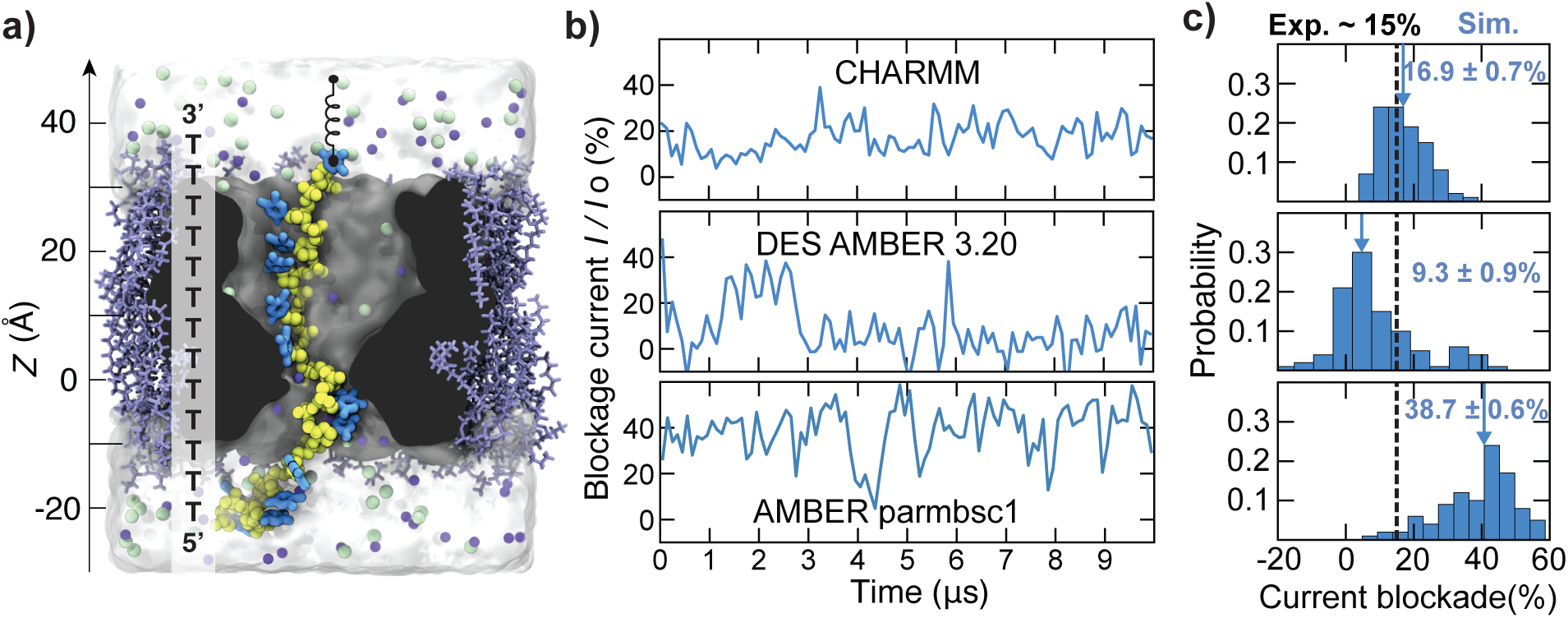
Blockade current of poly(dT) simulated using three popular MD force fields. **a,** Simulation system containing a poly(dT)_13_ strand (yellow for backbone, blue for bases) threaded through a truncated model of M1-MspA (gray), a lipid membrane (light blue), a water box (semi-transparent surface), K^+^ (purple) and Cl*^−^* ions (green). The C1*^′^* atom of the top DNA nucleotide (at the 3*^′^* end) is harmonically restrained to a point 30 Å above the nanopore constriction at the nanopore axis. **b,** Ionic current through the poly(dT)_13_-blocked nanopore, normalized by the open pore current (*I*_O_), in MD simulations carried out using the CHARMM36, DES-AMBER 3.20, and AMBER parmbsc1 parameter sets. Shown are 20 ps sampled currents averaged in 100 ns blocks. **c,** Histograms of the relative blockade currents. The histograms were constructed using 20 ns block-averaged data. The mean and the standard error of each histogram are shown in each plot. The dashed line and the solid arrow indicate the average experimental^21^ and simulated blockade value, respectively.

The simulation of the poly(dT) system using CHARMM36 produced the expected fluctuation of the ionic current, Fig. 4b. By splitting the current trace into 20 ns blocks, we constructed the histogram of the blockade current (*I*/*I*_0_), a quantity that can be readily compared to experiment. We find that the average current blockade, 16±0.7%, does not deviate much from the experimental value of ∼15%.^21^ Rather surprisingly, the simulations carried out using the DES-AMBER model significantly underestimated the current, whereas the AMBER parmbsc1 simulations overestimated the current by a wide margin, Fig. 4b,c. Thus, among the three force fields, CHARMM36 produced the closest quantitative agreement with experiment.

Visual inspection of the three simulations revealed strikingly different conformations of the poly(dT) strand at and near the MspA constriction, Fig. 5a. We quantified these conformations by plotting the center-of-mass *Z* coordinates of individual DNA bases over the course of the simulations, Fig. 5b, and characterizing their stacking propensity in, Fig. 5c, and near, Fig. 5d, the pore constriction. In the DES-AMBER simulation, the three nucleotides most proximal to the constriction adopted base-stacked conformations for the majority of the trajectory, blocking the passage of ions. An opposite effect was observed in the parmbcs1 simulations, where the DNA strand was observed to adhere to the nanopore surface, separating the bases of the two nucleotides most proximal to the constriction while forming stable multi-base stacks outside the constriction. This absence of bases in the MspA constriction increases its conductive volume and, hence, the nanopore current. In contrast, moderate base stacking was observed in the CHARMM36 simulation, resulting in current values in-between the two extreme cases of the AMBER force fields and matching the experimental value. Test simulations of poly(dC)_13_ threaded through a truncated M1-NNN MspA model in the same global orientation produced an average current blockade of 23.8±0.8% for CHARMM36 and 13.7±0.8% for DES-AMBER, SI Fig. 5, with the experimental value being ∼25%.^21^ Just like in the poly(dT) case, excessive base stacking was observed in the MspA constriction during the DES-AMBER simulation, causing a much deeper current blockade in comparison to the CHARMM36 simulation that showed moderate stacking of DNA bases.

**Figure 5:**
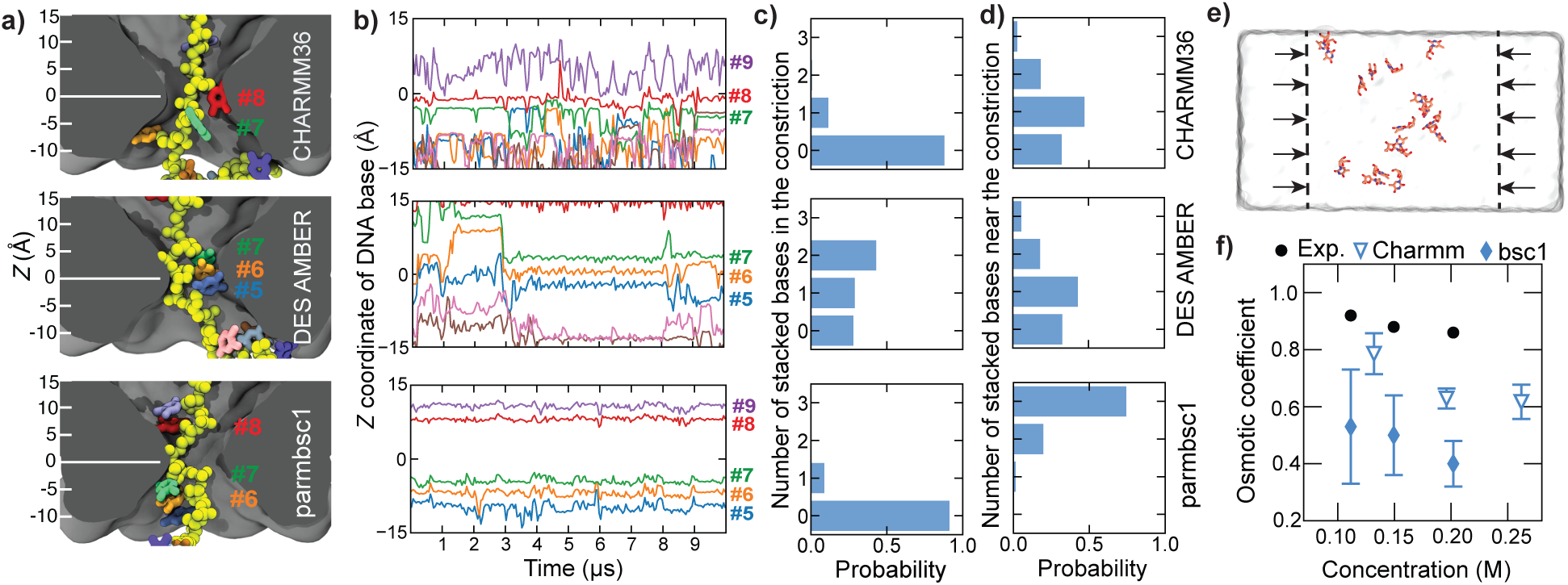
Force field affects the conformation of poly(dT) in MspA. **a,** Representative conformations of poly(dT) when simulated using the CHARMM36, DES-AMBER, or AMBER parmbsc1 force field. Nucleotide indices are indicated, with *Z* = 0 marking the pore constriction center. Residues are color-coded by index. **b,** *Z*-coordinate of select DNA bases *versus* simulation time, block-averaged in 50 ns. Residue indices are shown on the right; colors as in panel a. **c,** Probability of observing the specified number of stacked bases within an 8 Å slab centered at the nanopore constriction. Two bases were counted as stacked when their centers of mass were within 5 Å of one another and their interplanar angle less than 45*^◦^*. Only neighboring bases were considered for this analysis. **d,** Probability of observing the specified number of stacked bases in the near-constriction region defined as a 30 Å slab centered at the nanopore constriction but excluding the central 8 Å slab. **e,** Setup of osmotic pressure simulations. Deoxythymidine nucleosides are confined to the internal volume via two half-harmonic potentials (dashed lines). The osmotic pressure is computed as the average applied force divided by the system’s cross section area. **f,** Osmotic coefficients of thymine solutions from CHARMM36 and AMBER parmbsc1 simulations and experiment. The experimental and parmbsc1 data are from Ref. 92. Each CHARMM data point represents an average of three 70 ns replica simulations; the error bars indicate the standard deviation among the replicas.

To further probe the accuracy of base stacking interactions, we measured the osmotic pressure of a deoxythymidine solution as a function of its concentration, an approach commonly used to refine non-bonded parameters.^51,67^ In these simulations, thymine nucleosides were confined to compartment of pre-defined concentration using two half-harmonic potential, Fig. 5e. The the osmotic pressure was inferred from the average force exerted by the potential on the nucleotides. If stacking interactions are overly strong, the nucleotides would aggregate, reducing the osmotic pressure. The CHARMM36 simulations produced osmotic coefficients that were more consistent with the experimental values than those previously reported^92^ for the parmbsc1 model, Fig. 5f. The latter study found the parmbsc1 model to consistently underestimate the osmotic pressure due to excessive nucleosides clustering. ^92^

Taken together, our test simulations underscore the sensitivity of the blockade current to the force field model used for the simulations and emphasize the need for accurate description of base stacking interactions. Among the three force field tested, the CHARMM36 force field is found to provide the best overall agreement with experiment, despite mild overestimation of base stacking interactions.

### Refinement for DNA–protein interactions

In the previous section, CHARMM36 was identified as the most reliable force field for the simulation of a blockade current caused by the presence of ssDNA in a biological nanopore. We found this force field model, however, inadequate to reproduce the blockade current produced by a poly(dA) strand threaded through MspA in the 3*^′^*-*trans* orientation,^34^ Fig. 6a. In Fig. 6b, we reproduce the ionic current trace from that simulation. The ionic current is seen to drop and remain nearly zero for the last ∼3 *µ*s of the simulation. Corresponding to the fully blocked current state, the DNA adopts a persistent, highly stacked conformation, adhering to the inner surface of the pore, Fig. 6c (bottom). In contrast, parts of the same MD trajectory that correspond to a blockade level consistent with experiment (∼25%) feature much less compact DNA configurations, Fig. 6c (top).

**Figure 6:**
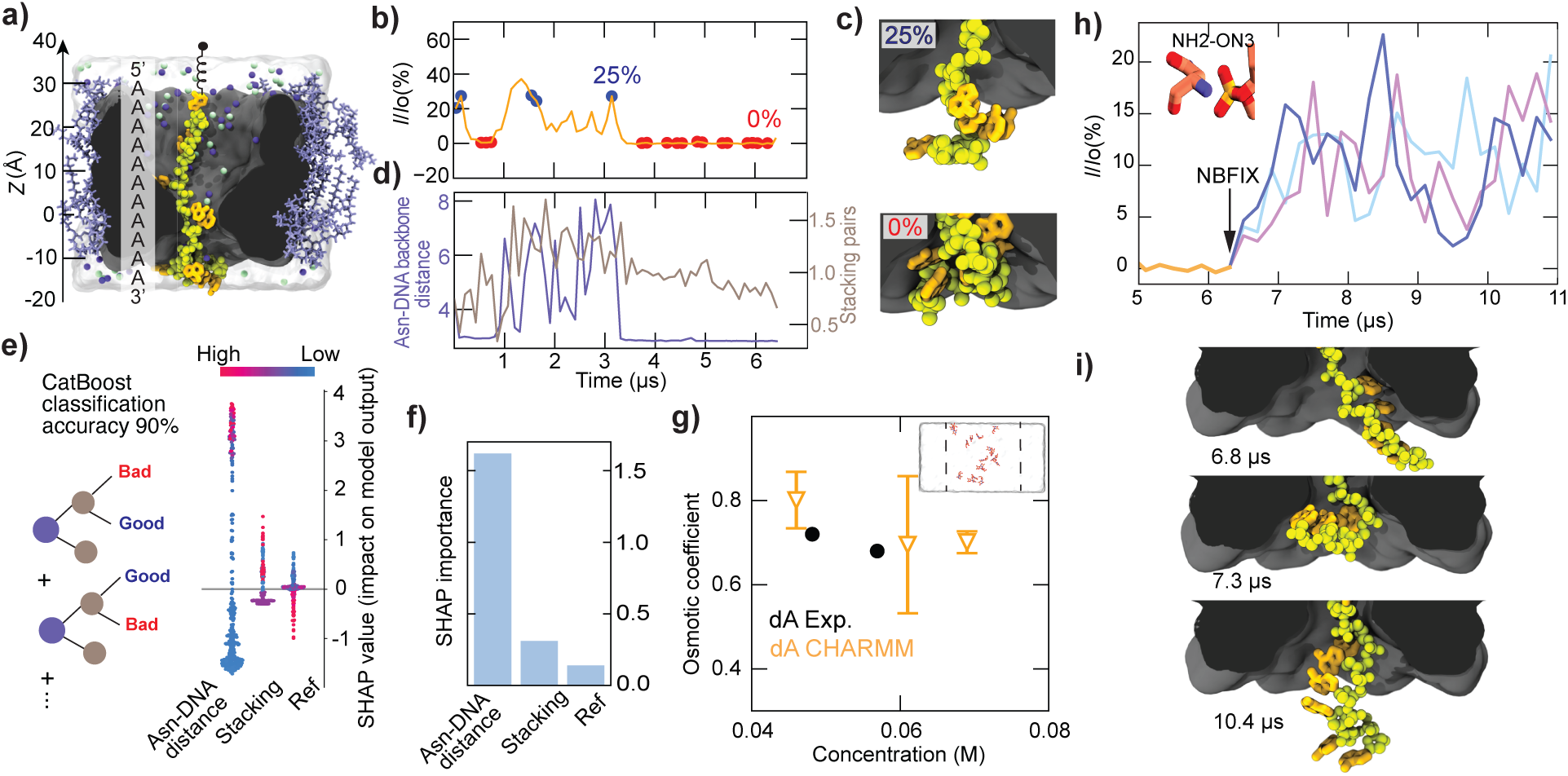
CHARMM36 overestimates the strength of specific DNA–protein interactions. **a,** Simulation system containing a poly(dA)_11_ strand threaded through truncated M1 MspA in the 3*^′^*-*trans* orientation. **b,** Simulated ionic current blockade (*I/I*_O_) *versus* time. The blue and red points mark instances when the simulated blockade current matches the experimental value (∼25%) or corresponds to a nearly complete blockade. Adapted from Ref. 34; © 2016 American Chemical Society. **c,** Representative conformations of poly(dA) corresponding to an experimental blockade current (top) or a nearly complete current blockade (bottom). **d,** Nearest distance between a non-hydrogen atom of an Asn_93_ MspA residue and the DNA backbone (purple, left axis) and the number of stacked bases among the four nucleotides nearest to the 3*^′^* end of the DNA (gray, right axis) *versus* simulation time. The plots show 100 ns block average of 100-ps sampled data. **e,** (left) Schematic of a CatBoost classifier used to distinguish “good” (20–30% blockade) from “bad” (*<*0.5% blockade) conformations using as features the number of stacked bases, the Asn_93_-DNA backbone contact distance, and one lipid’s phosphate *Z* coordinate (a reference feature). (right) SHAP analysis of model outputs, where red-to-blue scale indicates features of high-to-low impact on blockade current prediction. **f,** Mean absolute SHAP values ranking feature importance. **g,** Osmotic coefficients of deoxyadenosine solution as a function of its concentration. Experimental data ^92^ (black) are compared to CHARMM36 simulations (orange). The inset illustrates the simulation setup. Each CHARMM36 data point represent an average over three 50 ns replica simulations and the error bars denote the standard deviation among the replicas. **h,** Continuation of the simulation featured in panel b after applying the NBFIX correction to all Asn–DNA backbone interaction. The inset shows a close-up view of a hydrogen bond between Asn_93_ side-chain nitrogen (NH2) and a DNA backbone oxygen (ON3). Blockade current from three replica simulations are shown in different colors, each data point is an average of a 150 ns trajectory fragment. **i,** Representative conformations of poly(dA) from one replica simulation illustrating intermittent interactions between the DNA backbone and Asn_93_ in the presence of NBFIX corrections.

We hypothesized that this non-physical blockage arises from two potential artifacts: (i) overstabilized base stacking and (ii) overly strong interactions between the DNA and pore residues, particularly between residue Asn_93_ and the DNA backbone. To quantify these interactions, we tracked the number of stacking pairs near the constriction and the nearest distance between Asn_93_ and the DNA backbone over the simulation trajectory, Fig. 6d. The Asn_93_-DNA distance shows a strong correlation with the onset and persistence of the low-current conformations, suggesting it plays a dominant role. We thus hypothesized that the force field systematically overestimates the strength of Asn–DNA backbone interactions and that the nanopore confinement amplifies the effect of such overestimation.

To further validate the relative importance of these features, we combined our previously developed Steric Exclusion Model (SEM)^72^ of nanopore current with SHapley Additive ex-Planations (SHAP) ^74^ to evaluate the contribution of various structural descriptors to the blockade current value. Three thousands and eight hundreds instantaneous system’s configurations were extracted from the 6 *µ*s trajectory and labeled to contain “good” or “bad” conformations based on the agreement between the SEM-predicted and experimentally observed currents. Note that using SEM greatly reduced the error of instantaneous current determination without compromising its accuracy, as shown in our previous work.^72^ Each conformation was characterized by three features: (i) the number of stacking pairs among the four nucleotides below the constriction, (ii) the nearest distance between Asn_93_ and the DNA backbone, and (iii) a reference feature, which we chose to be the *Z*-coordinate of a lipid phosphate atom. These features were used to train a CatBoost classifier, ^73^ which achieved 90% classification accuracy, Fig. 6d. Our SHAP analysis revealed that the Asn_93_–DNA interaction contribute most significantly to misclassified conformations with a much higher importance score than the stacking or the reference features, Fig. 6e,f.

To independently assess the role of stacking, we conducted osmotic pressure simulations using solutions of deoxyadenosine monomers. CHARMM-based simulations showed good agreement with experimental osmotic coefficients, Fig. 6g, indicating that base stacking is not significantly overestimated in this force field model, Fig. 6f. These findings support the conclusion that the dominant factor driving abnormal current blockades in poly(dA) simulations is the excessive interaction between the pore residues and the DNA, rather than base stacking.

Further examination revealed that the aberrant stickiness of the DNA to the MspA pore in our CHARMM36 simulations primarily stems from strong interactions between the oxygen atoms of the DNA backbone and the nitrogen atoms of the carboxamide group of asparagines, Fig. 6h (inset). To reduce the strength of these interactions, we increased the *σ* parameter of the corresponding Lennard-Jones potential by 4% via a custom NBFIX correction. Specifically, the correction applied to the following three O–N interaction pairs: the nitrogen atom of any asparagine (NH2) group and the phosphate oxygens on DNA backbone (ON2) and (ON3) and one sugar oxygen (ON6). Table 2 lists the parameters that were affected by our correction. We then resumed the simulations of the poly(dA) system from the endpoint of the prior trajectory. The three replica simulations showed a consistent increase of the ionic current and disappearance of the fully blocked state, Fig. 6f,i. Importantely, the use of the same NBFIX corrections in CHARMM36 simulation of the poly(dT) system did not produce noticeable deviations of the blockade current from the experiment value, SI Fig. 6.

**Table 2:** NBFIX for asparagine–DNA backbone interactions.

|  |
| --- |
| NH2–ON2 |
| $R_{\min} = 3.76 \text{ \AA}, \quad \epsilon_{\min} = -0.17 \text{ kcal/mol}$ |
| NH2–ON3 |
| $R_{\min} = 3.69 \text{ \AA}, \quad \epsilon_{\min} = -0.15 \text{ kcal/mol}$ |
| NH2–ON6 |
| $R_{\min} = 3.76 \text{ \AA}, \quad \epsilon_{\min} = -0.17 \text{ kcal/mol}$ |

### Validation of the NBFIX corrections

We experimentally assessed the impact of the NBFIX corrections by measuring the residual current blockade under 150 mV bias using four single-stranded poly(dT)_40_ probes that contained abasic sites (nucleotides lacking bases) at four positions: nucleotides #12–15, #13–16, #14–17, or #12–17, Fig. 7a. Using such abasic constructs allowed us to directly probe the interaction of the DNA backbone and the asparagine side chains at the nanopore constriction, eliminating possible concerns regarding specific interactions between the nucleotide bases and their stacking. Each probe was modified with a biotin moiety at the 5*^′^* terminus and bound to streptavidin, which served as a physical stopper during nanopore translocation. When electrophoretically driven through the MspA nanopore, the streptavidin prevented full translocation, immobilizing the DNA within the pore. The top panel of Fig. 7b shows a typical ionic current trace experimentally recorded from the specified abasic construct. Among the four probes, the one with abasic sites at positions 12–17 exhibited the highest residual current, followed by the 13–16 construct, which residual current was only 1.5% less, Fig. 7c. This observation is consistent with previous reports indicating that nucleotides occupying positions 13–16 reside within the MspA constriction zone and dominate the ionic current readout and that nucleotides flanking the sensing region do not contribute significantly to current blockage.^21^ Blockade current was also measured for the 13–16 abasic construct under 180 mV, which showed a considerable increase of the relative blockade current compared to the identical measurement carried out at 150 mV, Fig. 7c. We hypothesized that stretching of the DNA backbone within the MspA constriction is responsible for this effect.

**Figure 7:**
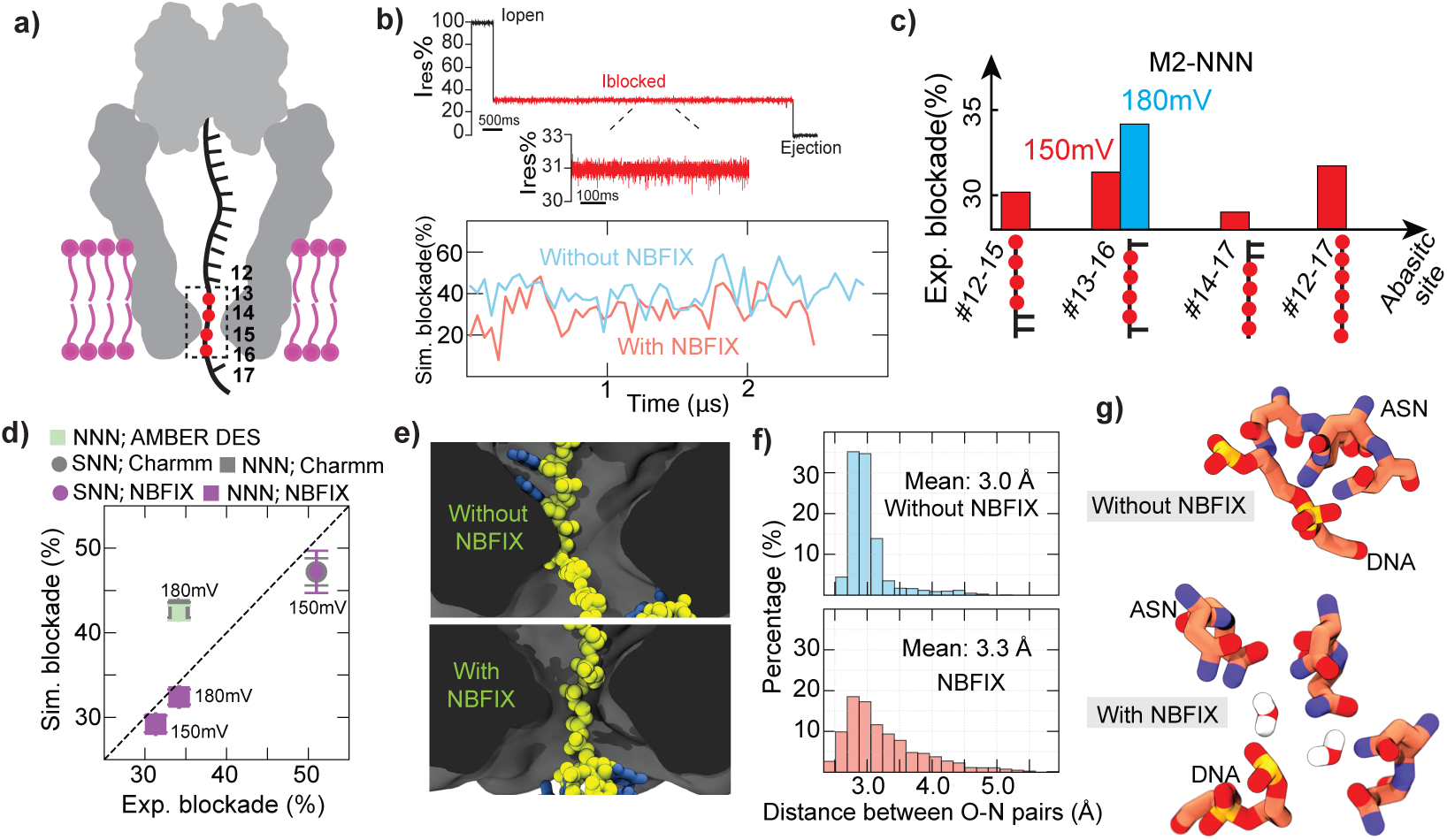
Experimental validation of the DNA backbone–asparagine NBFIX. **a,** Schematic of the experimental setup for measuring current blockades from abasic-site constructs. Streptavidin immobilizes the ssDNA within MspA, placing the abasic sites near the constriction. Nucleotides #13–16 (boxed) correspond to the primary sensing region. **b,** (Top) Representative current recording from poly(dT)_40_ containing abasic sites at positions #13-16, bound to streptavidin and being trapped in M2-NNN at +150 mV and ejected at −150 mV. (Bottom) Blockade current from a dT_4_(iSdp)_4_T_4_ construct threaded 3*^′^*-end *trans* side through the truncated model of M2-NNN MspA, when simulated with and without the Asn–DNA backbone NBFIX. Data show 50 ns block averages of 100 ps sampled current. **c,** Experimentally measured relative blockade current for oligonucleotides containing abasic nucleotides at different positions. Measurements at 150/180 mV are shown using red/blue bars. **d,** Experimental *versus* simulated relative blockade currents for the #13–16 abasic construct in M2-NNN and M2-SNN MspA. **e,** Representative conformations of the abasic construct within the constriction of M2-NNN MspA, simulated with (bottom) or without (top) the NFBIX corrections. **f,** Distribution of the closest distance between any oxygen atoms on the abasic site and any NH2 atom of Asn_90_ and Asn_91_ in the simulations carried out with (bottom; red) or without (top; blue) the NBFIX corrections. The distance was sampled every 100 ps over the ∼2.5 *µ*s trajectories. **g,** Close-contact configurations between the abasic DNA construct and Asn residues at the MspA constriction, illustrating conformations that are sampled less frequently after applying our NBFIX corrections (bottom) than in the standard CHARMM (top).

Matching these experiments, we conducted all-atom MD simulations of the abasic constructs using the following three force field models: the standard CHARMM36 force field supplemented by the CUFIX corrections, the standard DES-AMBER model^38^ and the CHARMM36 force field supplemented by both the CUFIX and the Asn-DNA NBFIX corrections. The simulation systems were constructed starting from a pre-equilibrated configuration of a poly(dT)_13_ strand threaded through a truncated M2-NNN MspA nanopore. The dT_4_(idSp)_4_dT_4_ strand was constructed by mutating four consecutive thymines near the MspA constriction into abasic sites, which was expected to match the placement of the #13–16 abasic residues in the corresponding experiment. As we found the relative blockade current to sensitively depend on the voltage applied in experiment, we took extra care to match the voltage applied in experiment to the one applied in the simulation. Because the truncated model of MspA lacks the access resistance of the full-scale nanopore system, the nominal transmembrane bias applied in the simulations may not match exactly the bias applied in experiment. To match the two, we simulated the open pore current through the truncated model of MspA under several values of the applied voltage. Following that, we determined the nominal values of the applied bias that produced, in simulations, the same scaled open pore current as in experiments carried out under 150 and 180 mV, SI Fig. 7. The systems were then simulated for ∼2.8 *µ*s using the three force field models under the transmembrane bias determined according to the above current matching procedure, SI Table 7.

The blockade current obtained using the standard CHARMM36 parameters and the CUFIX corrections, Fig. 7b, reached ∼43% at a bias matching the 180 mV experimental condition, overestimating the experimental value by approximately 25%, Fig. 7d. SHAP analysis reveals that excessive DNA–protein interactions contribute the most to such overestimation, SI Fig. 8. Equivalent simulations performed using DES-AMBER and the TIP4P-D water,^65^ a combination that has previously found to improved description of ssDNA–protein interactions,^38^ yielded relative blockade values that overlapped with the standard CHARMM result, Fig. 7d, similarly overestimating the simulated current over the experimental data.

Introducing the NBFIX corrections to Asn–DNA backbone interactions substantially improved the agreement between simulation and experiment. The simulated relative blockade decreased from 43% to 32.5%, aligning closely with the experimental value at 180 mV, Fig. 7b,d. When simulations were repeated under a lower transmembrane voltage, 150 mV, the NBFIX-refined model again reproduced the experimentally observed voltage-dependent decrease of the blockade current, demonstrating that the refinement captures not only the magnitude but also the voltage dependence of the ionic current. Further analysis showed that the average distance among the neighboring phosphates of the abasic segment increased by 0.02 Å, on average, along the pore axis when the voltage was increased from 150 to 180 mV.

To test robustness of the correction, we repeated both simulations and experiments using the M2-SNN mutant and the same abasic construct (#13–16). For this pore variant, the standard CHARMM36 / CUFIX matched experiment more closely, Fig. 7d, supporting our assessment that the large deviations observed for M2-NNN originate from specific interaction between DNA backbone and the Asn residues of the M2-NNN constriction. Importantly, applying the NBFIX correction in the SNN simulations maintained the quantitative agreement between experiment and simulations, Fig. 7d, exemplifying the amino acid-type specificity of the correction.

Visual inspection of the simulation trajectories revealed that the standard CHARMM force field favored excessive binding between the DNA backbone and the pore, Fig. 7e top, which increases the volume accessible for the ion flow and, hence, the current. In contrast, the NBFIX-corrected simulations reduced this binding, Fig. 7e bottom, lowering the conductive volume and resulting in blockade currents that matched experiment. We quantified this effect directly by computing the distributions of the nearest Asn–DNA backbone distance, *i.e.*, the nearest distance among all atom pairs involving the NH2 atom of Asn_90_ or Asn_91_ and either ON2, ON3, or ON6 atoms of the DNA backbone. The corrections notably reduced the frequency of short-range contacts while increasing the mean distance by 10%, Fig. 7f. Similar to the action of other NBFIX corrections,^53^ the Asn–DNA backbone NBFIX allowed water molecules more frequently mediate the asparagine–backbone interaction, favoring out direct waterless contacts, Fig. 7g.

Previous studies ^21^ and our own work, Fig. 7c, indicate that nucleotides #13–16 of a DNA strand must reside within the sensing region of MspA when the biotinylated ssDNA is trapped within MspA using a NeutrAvidin anchor, Fig. 7a. Thus, we examined whether the NBFIX corrections influence the DNA positioning relative to the sensing region. Our simulation system consisted of a full-length M2-NNN MspA, a poly(dT)_21_ strand partially threaded through the MspA constriction, and a NeutrAvidin tetramer bound to the 5*^′^* biotinylated end of the DNA, Fig. 8a. Note the reversal of the global DNA strand orientation in comparison to the systems examined previously in Fig. 4 and SI Fig. 6. This system was simulated for ∼ 2 *µ*s using the CHARMM36, the CUFIX corrections, and with or without the Asn–DNA backbone NBFIX.

**Figure 8:**
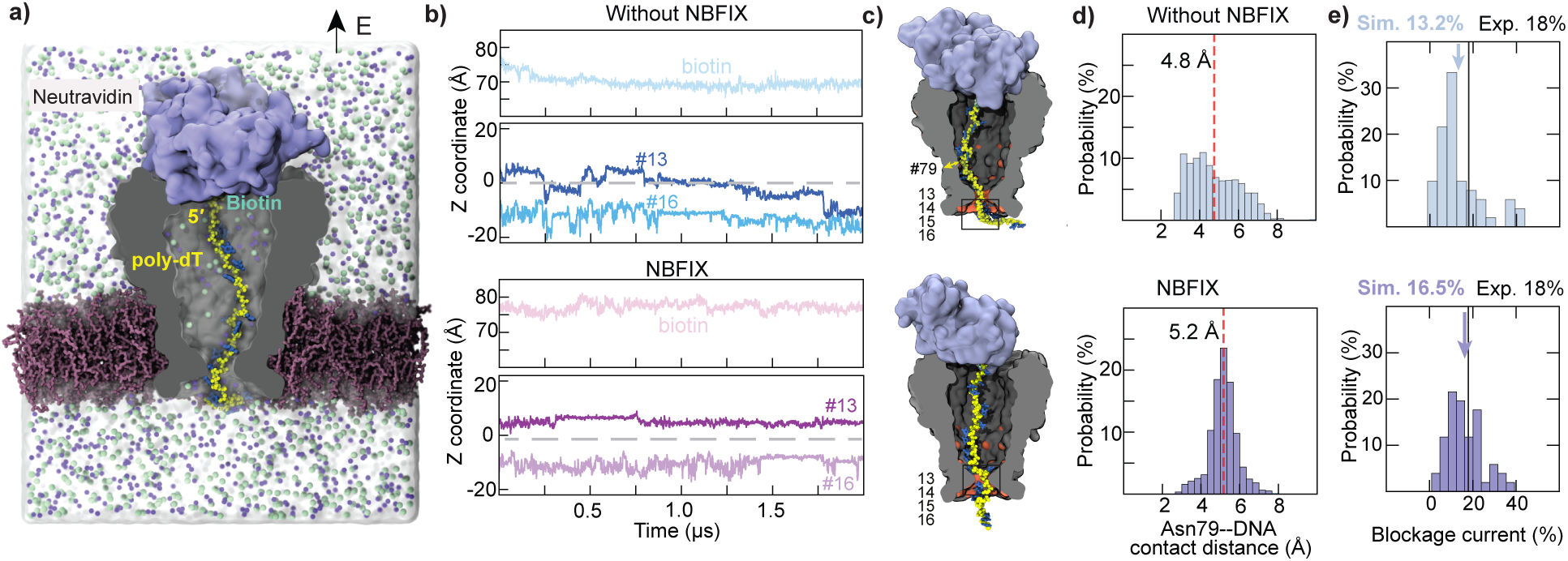
Asn–DNA backbone NBFIX improves description of full-length MspA system. **a,** Full-length M2-NNN MspA system containing a poly(dT)_21_ strand tethered via its 5*^′^* end to a biotin that is bound to NeutrAvidin. **b,** CoM *z*-coordinate of the biotin, thymine #13 and #16 nucleotides over the 2 *µ*s Anton 3 simulations carried out with (bottom) or without (top) the Asn–DNA backbone NBFIX. Dashed horizontal lines indicate the MspA constriction (CoM of residues #90 and #91). **c,** Snapshots illustrating the representative configurations (at 1.5 *µ*s) of the simulated systems. Thymine positions #13–16 are enclosed by a black box outline. The yellow arrow indicates the interaction between the DNA backbone and Asn_79_. Orange patches indicate the locations of Asn side chains. **c,** Distributions of thymine #13 and #16 *z* coordinates. Dashed lines mark the MspA constriction. **d,** Contact-distance distributions between the NH_2_ atom of MspA residue Asn_79_ and the DNA backbone atoms, comparing simulations carried out with (bottom) or without (top) the NBFIX. The peak shifts to larger distances when NBFIX is applied, indicating weakened interaction and reduced DNA stretching. Average values are provided in the panels. In panels b–c, data were collected every 1.2 ns. **e,** Distribution of the relative blockade current collected over the final 1 *µ*s of the respective trajectories. Instantaneous current values were averaged in 20 ns blocks. NBFIX shifts the mean blockade (arrows) from 13.2% to 16.5%. The experimental value of 18% is indicated by a solid lines. Note the reversal of the strand direction relative to the system featured in Fig. 4 and hence the difference in the experimental current value.

We find that introducing NBFIX markedly improves the consistency between simulations and experiments by placing the expected fragment of the DNA strand within the sensing region of the nanopore. Without NBFIX, the nucleotides #13–16 frequently shift below the constriction, spending most of the trajectory outside the experimental sensing zone, Fig. 8b (top). In contrast, these nucleotides remained near the pore constriction throughout the simulation that employed the Asn–DNA backbone NBFIX, Fig. 8b (bottom).

Inspection of the simulation trajectories reveals the origin of this improvement. In the absence of NBFIX, NeutrAvidin is gradually pulled downward into the lumen of MspA due to attractive interactions between the DNA backbone and the protein surface, particularly a cluster of asparagine residues in the lower pore region, Fig. 8c (top). The NBFIX correction alleviates this excessive attraction, preventing the anchor protein from being dragged into the pore and allowing NeutrAvidin to remain more relaxed outside the vestibule, Fig. 8c (bottom). This reduction of spurious protein–DNA interaction is also evident at the atomic level: the contact between Asn_79_ at the inner MspA surface away from the constriction and the DNA backbone is significantly weakened upon applying the NBFIX corrections, increasing the Asn_79_–DNA backbone contact distance, Fig. 8d. Consistent with these structural changes, the current blockade calculated from the MD trajectories of the full-length system is also affected by the introduction of NBFIX. The simulated blockade current with NBFIX is 16.5%, in closer agreement with the experimental value of 18% compared to 13.2% observed without NBFIX, Fig. 8e.

## CONCLUSIONS

In this work, we systematically investigated MD protocols for all-atom simulation of ion transport through biological nanopore. We showed that a properly calibrated and carefully setup all-atom MD simulation can accurately reproduce the ionic current through biological nanopore MspA and its mutant variants. Further, the same simulation can accurately characterize the selectivity of ion transport Our study has also identified several factors that may lead to an apparent disagreement between experimental and simulated open-pore current values. Chief among them is the systematic overestimation of the bulk ion conductivity by the MD force field, which is easy to correct by scaling the simulated ionic current with the ratio of experiment and simulated bulk electrolyte conductivity. A more subtle yet rather common problem is having the steady-state bulk ion concentration that is different from the experimental one. As we show in this work, such an inaccurate system setup can originate from using standard tool to solvate and ionize the system. That is, while such tool place water and ions in the correct ratio, subsequent redistribution of the water and ions within the simulated system can either increase or decrease ion concentration in the bulk regions of the systems, causing the ionic current simulations to be conducted at an effectively different bulk ion concentration. We suggest to carefully examine the steady state ion concentration in the bulk regions of the system and adjust the concentration by either adding or removing ions. MD simulations of MspA are particularly prone to this effect because of the high internal volume of this nanopore. An alternative post-simulation remedy is to scale the simulated currents with the ratio of experiment and simulation bulk conductivities using the simulated bulk conductivity value at the bulk concentration actually realized in the simulation.

Another cause for an apparent disagreement between simulated and experiment open pore current may stem from the difference in the access resistance. In experiment, the access resistance is well approximate by a semi-spherical volume of bulk electrolyte on each side on the nanopore. In an MD simulation, the access resistance volume is rectangular and, most importantly, has a non-trivial dependence on the system size because of the periodic boundary conditions. Setting up a system to have a golden ratio of its dimensions^82^ is a good first approximation, but further adjustment are likely needed to account for asymmetric shape of the nanopore. One should be careful when comparing simulated ion selectivity to reverse potential measurement. The first examines ions selectivity under symmetric buffer conditions, arguable the most relevant conditions for nanopore experiment, whereas the second inherently requires asymmetric buffer leading to the experimental selectivity measurement to depend on the polarity and the magnitude of the concentration gradient. As we showed in this work, good agreement between simulation and experiment is obtained when the simulations are performed under asymmetric buffer conditions. Finally, the choice of force field can affect the open pore conductance. In this work, we found the combination of CHARMM36M, TIP3P water and CUFIX^53^ corrections to ion–ion and ion–pore interactions to provide the optimal agreement with experiment.

Our assessment of three off-the-shelf force fields for reproducing blockade current from DNA homopolymers revealed that none of them had the required accuracy. Surprisingly, we found the older CHARMM force field to outperform more recent AMBER variants likely because of its more accurate treatment of base stacking, as confirmed by osmotic-pressure benchmarks. Motivated by these observations, we sought to improve CHARMM36 by focusing on refinement of DNA-protein interactions at the nanopore constriction. We identified a key artifact that, in prior simulations,^34^ led to DNA strands becoming abnormally compact and adhering to the pore wall, completely blocking the ionic current in a major contradiction to experiment.^21^ Our analysis revealed that overly strong interactions between the DNA backbone and specific pore residues– asparagines–were responsible for this artifact. To correct for this artifact, we introduced surgical NBFIX corrections to specific non-bonded Lennard-Jones parameters describing DNA–pore interactions. The refined model successfully reduced spurious adhesion of the DNA backbone to asparagines, unclogging the nanopore. To validate these refinements, we designed and conducted nanopore experiments using DNA strands that contained stretches of abacis residues, directly probing the DNA–pore interactions that were target of our force field refinement. Matching the setup of these experiments, additional MD simulations confirmed that our refined model accurately recapitulates the experimental current signatures. Importantly, our NBFIX corrections weakened only the problematic interaction without disrupting the original force field integrity.

While the progress described about is substantial, we do not yet have an all-atom MD force field capable of reproducing the DNA sequence specificity of ionic current blockades in biological nanopores. For example, while our refinement has improved the MD description of a poly(dA) blockade current, the simulated blockade is not yet in perfect agreement with experiment. Each apparent simulation–experiment discrepancy may arise from multiple sources, including electrolyte conductivity, ion–protein interactions, DNA–protein interactions, sampling limitations, or differences between the experimental and simulated systems. Analysis methods such as SHAP can identify dominant factors responsible for large discrepancies, when such factors are present, however, they are less effective when multiple small inaccuracies nearly equally contribute to the discrepancies. On a technical side, further progress is limited by the time scale required for an ssDNA to probe its microscopic configurations within the nanopore. Furthermore, while experiment has provided a sequence-to-current lookup table,^7^ such data are not ideal target for force field refinement as the measured current is averaged over the three to four nucleotides undergoing stochastic motion within the nanopore constriction. We intend to overcome this problem by designing custom DNA and RNA constructs that make one-to-one comparison between simulation and experiment possible.

Taken together, our results underscore the critical importance of accurate description of ion behavior and DNA-protein interactions for all-atom MD simulations of nanopore ionic current. By integrating improved physical models, targeted force field refinement, and experimental validation, we provide a robust framework for advancing the predictive capabilities of MD simulations for nanopore-based DNA, RNA and proteins sequencing and sensing applications. Beyond nanopores, we expect our refined parameterization of the MD force field to improve realism of all-atom MD simulations of a broad class of DNA–protein and RNA–protein systems.

## Acknowledgement

This work was supported by the grant from the National Institutes of Health (R21-HG011741 to M.C and A.A). The supercomputer time was provided through the ACCESS allocation MCA05S028 and the Leadership Resource Allocation MCB20012 on Frontera of the Texas Advanced Computing Center. We acknowledge computational resources provided by the Pittsburgh Supercomputing Center (PSC) on the Anton 3 and Anton 2 supercomputers. Anton 3 computer time was provided through NIH Grant 1R24GM154042, and Anton 2 computer time was provided through NIH Grant R01GM116961. These machines, which were made available by D. E. Shaw Research, enabled the long-timescale molecular dynamics simulations in this work. We thank Dr. Liang Xu, Dr. Bach Pham for his help during the initial stages of this project and Ms. Melika Shekari for her help with establishing a protocol for MD simulations under a gradient of ion concentration.

## Supporting Information Available

Supplementary tables detailing all MD runs; figures illustrating bulk conductivity simulations, system setup artifacts that automated setup tools may introduce, open pore simulations of MspA mutants using halide NBFIX, additional MD simulations of poly(dC) and poly(dT) systems, applied voltage calibration by matching open pore current, and SHAP analysis of abasic construct simulations. This information is available free of charge via the Internet at http://pubs.acs.org.

