## Supplementary Information for "Improving All-Atom Molecular Dynamics Models for Quantitative Prediction of Nanopore Blockade Current"

**SI Table 1.** MD simulations of open-pore M2-NNN MspA using CHARMM36 and CUFIX. The simulations investigated the effect of system size on the open-pore current. The standard errors were calculated by splitting each ∼400 ns trajectory into 50 blocks and treating the average within each block as an independent measurement.

| Bias | System size | Simulation length | Raw MD current | Scaled current |
|---|---|---|---|---|
| +180 mV | $65 \times 65 \times 80\,\text{Å}^3$ | 400 ns | $664 \pm 19$ pA | $405 \pm 12$ pA |
| | $110 \times 110 \times 120\,\text{Å}^3$ | 380 ns | $593 \pm 13$ pA | $361 \pm 8$ pA |
| | $160 \times 160 \times 200\,\text{Å}^3$ | 400 ns | $550 \pm 13$ pA | $335 \pm 8$ pA |
| | $111 \times 111 \times 233\,\text{Å}^3$ | 400 ns | $532 \pm 16$ pA | $324 \pm 10$ pA |

**SI Table 2.** MD simulations and experimental characterization of open-pore current through MspA mutants. The MD simulations were performed using the CHARMM36 force field and the CUFIX corrections; the system size was $110 \times 110 \times 120$ Å$^3$. The scaled MD current was obtained by multiplying the corresponding raw MD current by 0.61, which is the ratio of the experimental and simulated conductivity of 1 M KCl solution. The standard deviations were calculated by splitting each 270 ns trajectory into 50 blocks and treating the average within each block as an independent measurement.

| Bias | Mutant | Simulation length | Raw MD current (pA) | Scaled MD current (pA) | Experiment (pA) |
|---|---|---|---|---|---|
| +200 mV | GNS | 270 ns | $1067 \pm 64$ | $651 \pm 39$ | 734 |
| | GSN | | $1147 \pm 57$ | $700 \pm 35$ | 779 |
| | NGS | | $1104 \pm 28$ (restrained) | $673 \pm 17$ (restrained) | 797 |
| | NGS | | $1128 \pm 30$ (unrestrained) | $688 \pm 18$ (unrestrained) | 797 |
| | NNN | | $610 \pm 32$ | $372 \pm 20$ | 402 |
| | NNS | | $671 \pm 55$ | $409 \pm 34$ | 441 |
| | NSN | | $1091 \pm 37$ | $666 \pm 23$ | 665 |
| | SNN | | $879 \pm 33$ | $536 \pm 20$ | 600 |
| -200 mV | GNS | 270 ns | $-680 \pm 34$ | $-415 \pm 21$ | -500 |
| | GSN | | $-678 \pm 51$ | $-414 \pm 31$ | -545 |
| | NGS | | $-791 \pm 74$ (restrained) | $-483 \pm 45$ (restrained) | -506 |
| | NGS | | $-826 \pm 66$ (unrestrained) | $-504 \pm 40$ (unrestrained) | -506 |
| | NNN | | $-605 \pm 59$ | $-369 \pm 36$ | -506 |
| | NNS | | $-601 \pm 53$ | $-367 \pm 32$ | -505 |
| | NSN | | $-645 \pm 58$ | $-393 \pm 35$ | -560 |
| | SNN | | $-634 \pm 43$ | $-387 \pm 26$ | -535 |

**SI Table 3.** MD simulations and experimental characterization of open-pore current through MspA mutants. The MD simulations were performed using the CHARMM36 force field and the CUFIX corrections; the system size was $65 \times 65 \times 80$ Å$^3$. The scaled MD current was obtained by multiplying the corresponding raw MD current by 0.61, which is the ratio of the experimental and simulated conductivity of 1 M KCl solution. The standard deviations were calculated by splitting each 300 ns trajectory into 50 blocks and treating the average within each block as an independent measurement.

| Bias | Mutant | Simulation length | Raw MD current (pA) | Scaled MD current (pA) | Experiment (pA) |
|---|---|---|---|---|---|
| +200 mV | NAN | 300 ns | $1169 \pm 50$ | $713 \pm 31$ | 762 |
| | NLN | | $450 \pm 29$ | $275 \pm 18$ | 247 |
| | NTN | | $1067 \pm 45$ | $651 \pm 27$ | 701 |

**SI Table 4.** MD simulations of open-pore current through MspA mutants performed using CHARMM36, CUFIX and halide NBFIX of Orabi et al.[56] The scaled MD current is obtained by multiplying the corresponding raw MD current by 0.53, which corresponds to the ratio of the simulated conductivity of 1.14 M KCl electrolyte and experimental conductivity of 1 M KCl. See more details in SI Fig. S**??**. The standard deviations were calculated by splitting each 300 ns trajectory into 50 blocks and treating the average within each block as an independent measurement.

| Electrical bias | Mutant type | Simulation length | Restrained (pA) | Unrestrained (pA) | Scaled restrained (pA) | Scaled unrestrained (pA) | Experiment (pA) |
|---|---|---|---|---|---|---|---|
| +200 mV | GNS | 300 ns | $1100 \pm 33$ | $1168 \pm 35$ | $583 \pm 17$ | $619 \pm 19$ | 734 |
| | GSN | | $1247 \pm 42$ | $1289 \pm 48$ | $661 \pm 22$ | $683 \pm 25$ | 779 |
| | NGS | | $1226 \pm 37$ | $1265 \pm 37$ | $650 \pm 20$ | $670 \pm 20$ | 797 |
| | NNN | | $729 \pm 51$ | $808 \pm 25$ | $386 \pm 27$ | $428 \pm 13$ | 402 |
| | NNS | | $797 \pm 26$ | $847 \pm 29$ | $422 \pm 14$ | $449 \pm 15$ | 441 |
| | NSN | | $1194 \pm 52$ | $1014 \pm 31$ | $633 \pm 28$ | $537 \pm 16$ | 665 |
| | SNN | | $1003 \pm 44$ | $1219 \pm 48$ | $532 \pm 23$ | $646 \pm 25$ | 600 |
| -200 mV | GNS | 300 ns | $-739 \pm 50$ | $-749 \pm 52$ | $-392 \pm 27$ | $-397 \pm 28$ | -500 |
| | GSN | | $-781 \pm 76$ | $-881 \pm 51$ | $-414 \pm 40$ | $-467 \pm 27$ | -545 |
| | NGS | | $-873 \pm 24$ | $-866 \pm 27$ | $-463 \pm 13$ | $-459 \pm 14$ | -506 |
| | NNN | | $-687 \pm 41$ | $-806 \pm 57$ | $-364 \pm 22$ | $-427 \pm 30$ | -506 |
| | NNS | | $-683 \pm 40$ | $-737 \pm 22$ | $-362 \pm 21$ | $-391 \pm 12$ | -505 |
| | NSN | | $-765 \pm 39$ | $-813 \pm 45$ | $-405 \pm 21$ | $-431 \pm 24$ | -560 |
| | SNN | | $-738 \pm 85$ | $-836 \pm 89$ | $-391 \pm 45$ | $-443 \pm 47$ | -535 |

**SI Table 5.** Simulated and experimental ion selectivity of MspA mutants.

| Pore type | Bias | KCl concentration | NBFIX type | Simulation duration | Simulated ion selectivity | Experimental ion selectivity |
|---|---|---|---|---|---|---|
| M2–NNN | 180 mV | 1M (*cis*)/1M (*trans*) | CUFIX | 400 ns | 1.3 | — |
| | | | CUFIX + halide NBFIX | 300 ns | 2.2 | |
| | | | Standard CHARMM36 | 300 ns | 1.5 | |
| M2–NNN | Off | 3M (*cis*)/1M (*trans*) | CUFIX | 110 ns | 0.92 | 0.92 |
| | | | CUFIX + halide NBFIX | 110 ns | 1.6 | |
| M2–NGS | Off | 3M (*cis*)/1M (*trans*) | CUFIX | 110 ns | 1.44 | 1.42 |
| | | | CUFIX + halide NBFIX | 110 ns | 1.8 | |
| M2–NNS | Off | 3M (*cis*)/1M (*trans*) | CUFIX | 110 ns | 0.94 | 0.82 |
| | | | CUFIX + halide NBFIX | 110 ns | 1.3 | |

**SI Table 6.** MD simulations of poly(dT)$_{13}$ and poly(dC)$_{13}$ homopolymers on Anton 2 using three force fields models. The simulations were run at 180 mV.

| DNA sequence | Orientation | Force field | Simulation length | MD blockage | Exp. blockage[21] |
|---|---|---|---|---|---|
| $dT_{13}$ | 5′-*trans* | AMBER DES 3.20 | 10 $\mu$s | 9.3% | 15% |
| | | AMBER Parm bsc1 | 10 $\mu$s | 38.7% | |
| | | CHARMM36 | 10 $\mu$s | 16.9% | |
| $dC_{13}$ | 5′-*trans* | AMBER DES 3.20 | 10 $\mu$s | 13.7% | 25% |
| | | CHARMM36 | 4 $\mu$s | 23.5% | |

**SI Table 7.** Simulated blockade currents for poly(dA)$_{11}$ and abasic DNA constructs and three MspA mutants.

| **DNA sequence** | **Pore** | **Bias** | **Orientation** | **Blockage before NBFIX** | **Blockage after NBFIX** | **Exp. blockage** |
|---|---|---|---|---|---|---|
| $dA_{11}$ | M1–NNN | 180 mV | 3′-*trans* | 0% | 9.8% | 25% |
| $dT_4(idSp)_4T_4$ | M2–NNN | 180 mV | 3′-*trans* | 42.7% | 32.4% | 34% |
| | | | | AMBER DES: 42.1% | N/A | 34% |
| | | 150 mV | | *n/a* | 29.2% | 31% |
| | M2–SNN | 150 mV | | 47.2% | 47.2% | 51% |

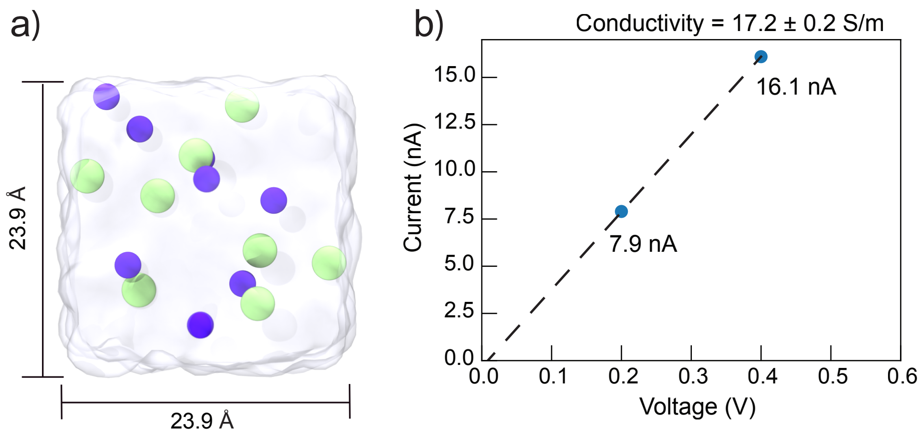


**SI Fig. 1. MD conductivity of 1 M KCl. a,** Cubic simulation system containing 1 M KCl. One side of the cubic unit cell $L = 23.9$ Å. Green and purple spheres represent $K^+$ and $Cl^-$ ions, respectively. **b,** Simulated ionic current under 200 mV and 400 mV bias. The bulk conductivity $\sigma = I/(V * L)$ is 17.2 $\pm$0.2 S/m, where the error estimated as $|\sigma_{400\mathrm{mV}} - \sigma_{200\mathrm{mV}}|/2$. The simulations were carried using the CHARMM36 parameter set, TIP3P water model and the CUFIX[51] corrections to $K^+ - Cl^-$ interactions. The temperature was maintained at 295 K using a Langevin thermostat with the dumping constant of 0.1 $ps^{-1}$ applied to all non-hydrogen atoms.

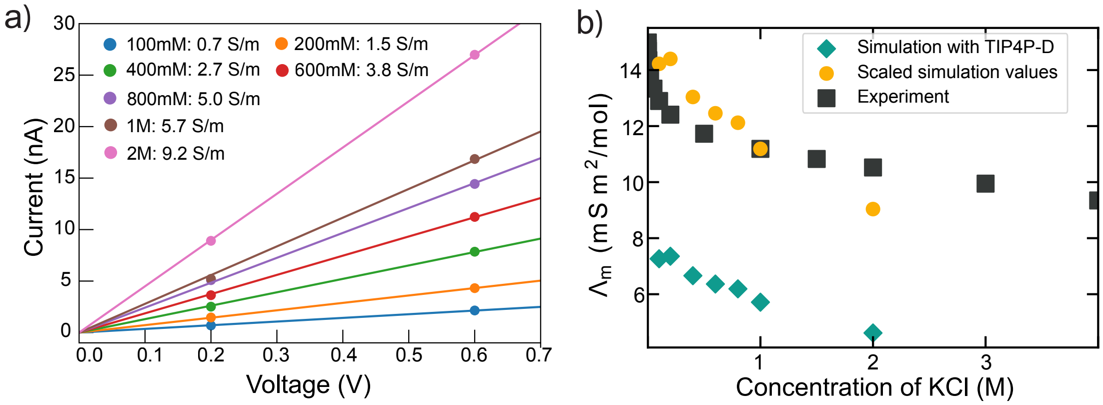


**SI Fig. 2. Bulk conductivity of KCl electrolyte in the TIP4P-D water model. a,** Current–voltage relationships obtained from MD simulations of bulk TIP4P-D[65] containing KCl at different concentrations. The simulations used the Joung–Cheatham ion parameters[64] without any NBFIX corrections. The electrolyte conductivity was computed as described in the caption to SI Fig. S**??**. All simulations employed a cubic unit cell with a side length $L = 48.8$ Å. Each system was equilibrated for 5 ns and simulated under applied electric field for 10 ns. **b,** Molar conductivity, $\Lambda_m$, as a function of KCl concentration. Green diamonds indicate raw simulation data obtained using the TIP4P-D water model. The black squares represent the experimental data. The orange circles denote the simulated conductivity values obtained after scaling all data points with the ratio of the experimental and simulation conductivities of 1M KCl solution.

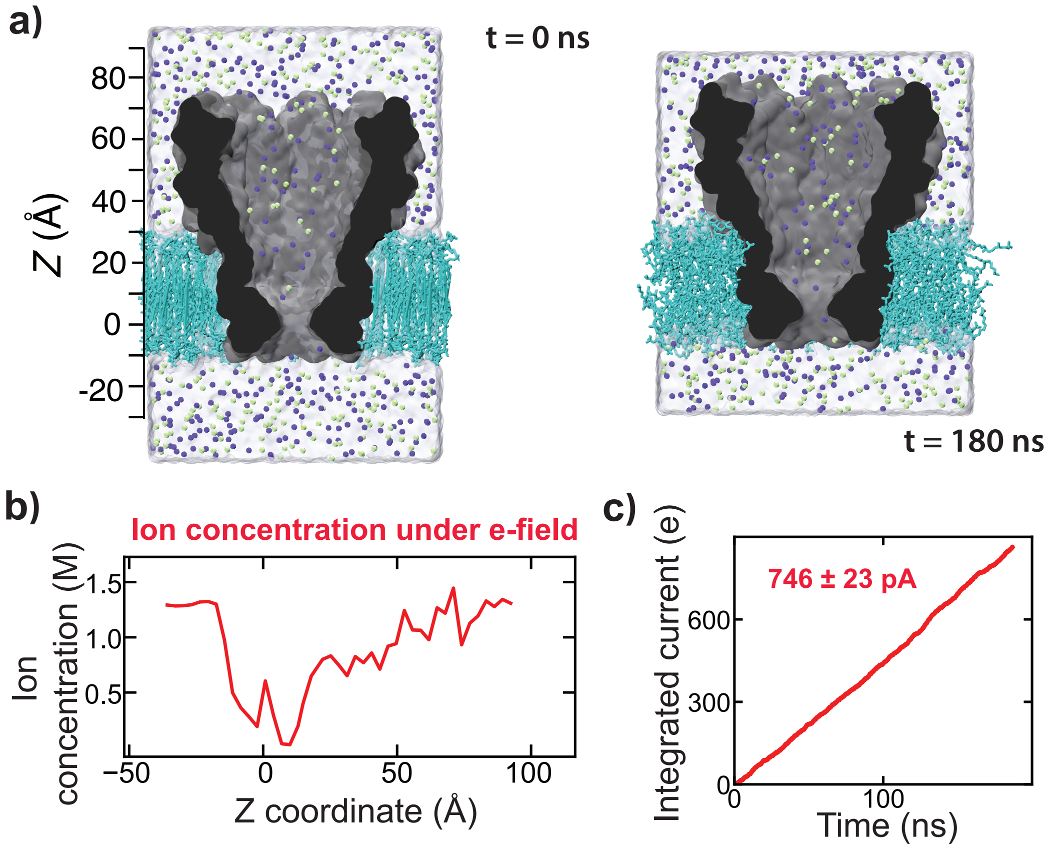


**SI Fig. 3. Automated setup tools can artificially increase bulk ion concentration. a,** All-atom model of M1 MspA as built by VMD (left) and at the end of a 180 ns applied electric field run (right). The solvated system was prepared by adding water to a dry membrane / MspA assembly using the `Solvate` plugin of VMD followed by the `Autoionize` plugin to add ions in the ion-to-water ratio corresponding to 1 M KCl concentration. During the 60 ns equilibration, the system's size changed as water solvated head groups of the lipid membrane. During the 180 ns simulation under a 180 mV bias, ions vacate the nanopore interior, increasing the ion concentration outside the nanopore to 1.25 M. **b,** Ion concentration profile along the nanopore ($z$) axis at the end of the applied electric field simulation. **c,** Total charge transported by ions over the course of the applied electric field simulation. The averaged simulated current (746 pA) scaled by the ratio of experimental and simulated conductivity of 1M KCl is 455 pA, which is 39% higher than the experimentally measured open pore current under similar conditions (328 pA).[21] Scaling the raw MD current with the ratio of experimental conductivity of 1M KCl and simulated conductivity of 1.25 M KCl (obtained by interpolation)—the steady-state bulk concentration observed during the applied electric field simulation—yields a current value of 364 pA, which is much closer to the experimental value.

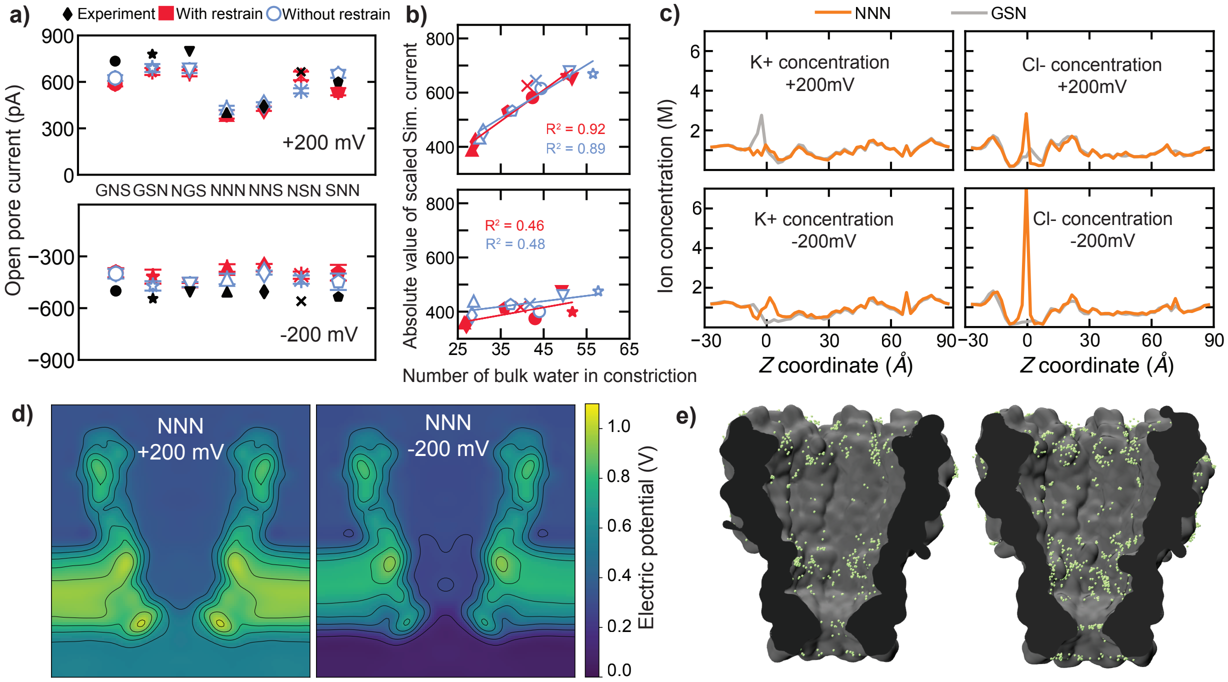


**SI Fig. 4. Simulated open pore current of MspA mutants obtained using CHARMM36 and both CUFIX and halide NBFIX corrections.** **a,** Open-pore current of seven MspA variants measured in experiment (black) and simulation (red/blue) under $\pm$200 mV. In half of the simulations (red), $C\alpha$ atoms of MspA were restrained to their crystallographic coordinates and other simulations (blue) were carried out in the absence of any residues. The simulated current values were scaled by a factor of 0.53 (current values shown in SI Table S**??**), which corresponds to the ratio of the simulated conductivity of 1.14 M KCl electrolyte and experimental conductivity of 1 M KCl. The simulated conductivity of 1.14 M KCl electrolyte was obtained by interpolation from the 1 M bulk value. We scaled the raw simulated currents using the 1.14 M simulated bulk conductivity (and not the 1 M value) because 1.14 M was the steady-state ion concentration established away from the nanopore in our ionic current simulations (see data in panel c). **b,** Magnitude of the simulated current *versus* number of bulk-like waters in the MspA constriction for simulation carried out with and without restrains. **c,** Trajectory-averaged concentration of $K^+$ and $Cl^-$ ions along the axis of the NNN and GSN pore. **d,** Electrostatic potential in M2 NNN MspA under $\pm$200 mV. The maps, averaged over the MD trajectory and the cylindrical symmetry of the nanopore, are shown for a plane encompassing the nanopore axis. **e,** Accumulated ions (multiple frames in VMD) within 2.5 Å of the M2-NNN MspA surface over 20 ns of the simulation (sampled per 0.24 ns) carried out using CUFIX (left) or CUFIX and halide NBFIX[56] (right).

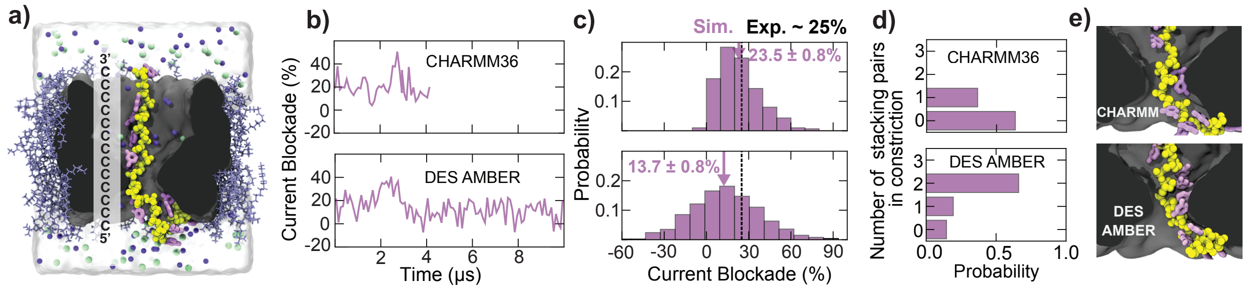


**SI Fig. 5. Relative blockade current of poly(dC) when simulated using CHARMM36 and AMBER-DES force fields. a,** Simulation system containing a poly(dC)$_{13}$ strand (yellow for backbone, pink for bases) threaded through a truncated model of M1-NNN MspA (gray), a lipid membrane (light blue), a water box (semi-transparent surface), $K^+$ (purple) and $Cl^-$ ions (green). The C1′ atom of the top DNA nucleotide (at the 3′ end of the strand) is harmonically restrained to a point above the nanopore along the nanopore axis. **b,** Histograms of the relative blockade currents. The histograms were constructed using 20 ns block-averaged data. The mean and the standard error of each histogram are shown in each plot. The dashed line and the solid arrow indicate the average experimental[21] and simulated blockade value, respectively. **c,** Probability of observing the specified number of stacked base pairs within the nanopore constriction in CHARMM36 (top) and AMBER-DES (bottom) simulations. **d,** Representative conformations of poly(dC))$_{13}$ when simulated using CHARMM36 (top) or AMBER-DES (bottom).

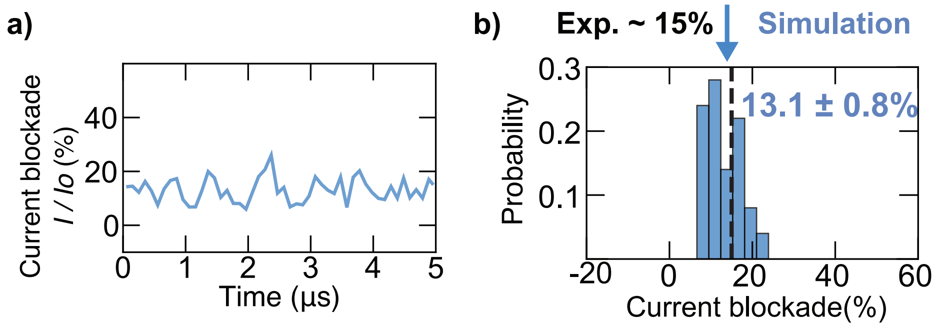


**SI Fig. 6. MD simulation of the poly(dT)$_{13}$ system using Asn–DNA phosphate NBFIX.** **a,** Ionic current through the poly(dT)-blocked nanopore, normalized by the open pore current ($I_O$), in MD simulations carried out using CHARMM36, CUFIX and the NBFIX between asparagine side chain nitrogen and the DNA backbone oxygens (Table 2 of the main text). Shown are 20 ps sampled currents averaged in 100 ns blocks. **b,** Histograms of the relative blockade currents. The histograms were constructed using 20 ns block-averaged data. The mean and the standard error of the histogram are shown. The vertical dashed line shows the average experimental blockade value[21] whereas the solid arrow indicates the simulated one.

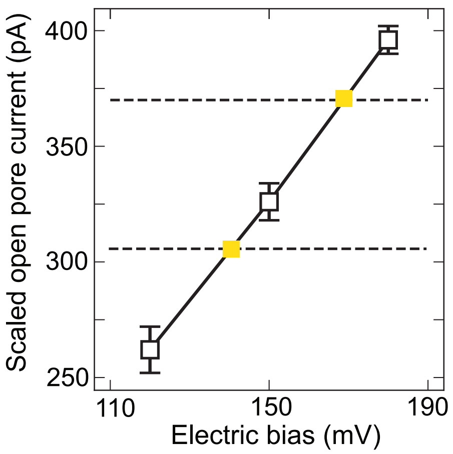


**SI Fig. 7. Choosing applied bias conditions by matching open pore currents.** Using the CHARMM36/CUFIX MD model, we determined the open pore current at 120, 150 and 180 mV. The simulations at 120 and 150 mV lasted ∼500 ns while the simulation at 180 mV lasted ∼1000 ns. The average values and the error bars were calculated by splitting each trajectory into 20 ns blocks and using the mean of each block as an independent data point. The current values were scaled using the ratio of the experimental and simulated bulk conductivity of 1 M KCl. The solid line represents a linear fit to the three points, passing through the origin. The horizontal dashed lines indicate the experimental open pore currents measured at 150 and 180 mV. By interpolation we find that, under 140 and 168 mV, the scaled simulated open pore current matches the experiment currents at 150 and 180 mV.

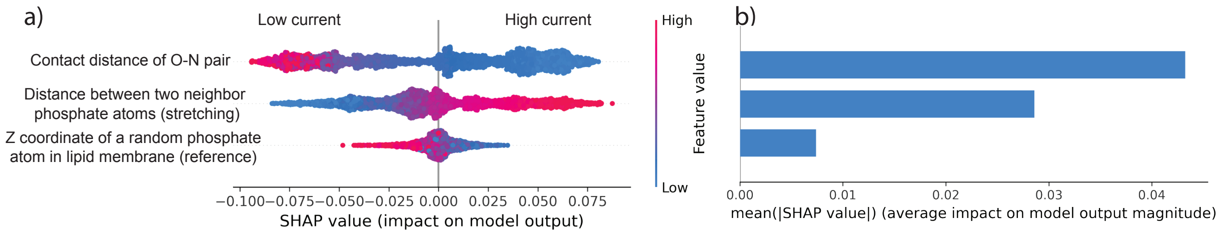


**SI Fig. 8. SHAP analysis of $dT_4(idSp)_4dT_4$ simulations performed using the CHARMM36 force field. a,** SHAP analysis of CatBoost classifier to distinguish "good" (32-36% blockade) from "bad" (>45% blockade) conformations using, as features, the shortest $Asn_{90,91}$–DNA backbone distance, the averaged phosphate-phosphate distance representing DNA stretching, and one lipid's phosphate $z$ coordinate (reference feature). **b,** Mean absolute SHAP values ranking feature importance, highlighting $Asn_{90,91}$–DNA backbone contacts as the dominant predictor.
